# Discovery of a Novel Inhibitor of Coronavirus 3CL Protease for the Potential Treatment of COVID-19

**DOI:** 10.1101/2020.09.12.293498

**Authors:** Britton Boras, Rhys M. Jones, Brandon J. Anson, Dan Arenson, Lisa Aschenbrenner, Malina A. Bakowski, Nathan Beutler, Joseph Binder, Emily Chen, Heather Eng, Holly Hammond, Jennifer Hammond, Robert E. Haupt, Robert Hoffman, Eugene P. Kadar, Rob Kania, Emi Kimoto, Melanie G. Kirkpatrick, Lorraine Lanyon, Emma K. Lendy, Jonathan R. Lillis, James Logue, Suman A. Luthra, Chunlong Ma, Stephen W. Mason, Marisa E. McGrath, Stephen Noell, R. Scott Obach, Matthew N. O’Brien, Rebecca O’Connor, Kevin Ogilvie, Dafydd Owen, Martin Pettersson, Matthew R Reese, Thomas F. Rogers, Michelle I. Rossulek, Jean G. Sathish, Norimitsu Shirai, Claire Steppan, Martyn Ticehurst, Lawrence W. Updyke, Stuart Weston, Yuao Zhu, Jun Wang, Arnab K. Chatterjee, Andrew D. Mesecar, Matthew B. Frieman, Annaliesa S. Anderson, Charlotte Allerton

## Abstract

COVID-19 caused by the SARS-CoV-2 virus has become a global pandemic. 3CL protease is a virally encoded protein that is essential across a broad spectrum of coronaviruses with no close human analogs. The designed phosphate prodrug PF-07304814 is metabolized to PF-00835321 which is a potent inhibitor *in vitro* of the coronavirus family 3CL pro, with selectivity over human host protease targets. Furthermore, PF-00835231 exhibits potent *in vitro* antiviral activity against SARS-CoV-2 as a single agent and it is additive/synergistic in combination with remdesivir. We present the ADME, safety, *in vitro*, and *in vivo* antiviral activity data that supports the clinical evaluation of this compound as a potential COVID-19 treatment.

## Introduction

In December 2019, COVID-19 was identified as a new, potentially fatal, respiratory infection caused by severe acute respiratory syndrome coronavirus 2 (SARS-CoV-2)^1,2^. Unlike previous coronavirus outbreaks that receded relatively quickly, the resultant COVID-19 pandemic spread across the globe. As of December 2020, over 75 million people have been infected and over 1.6 million people have died globally with no approved drugs available to treat the disease^3^.

The RNA-dependent RNA polymerase (RdRp) inhibitor remdesivir is currently undergoing clinical investigation for the treatment of SARS-CoV-2 and was granted emergency use authorization by the U.S. Food and Drug Administration (FDA) in May 2020^4^. To date, the results of trials of remdesivir have been mixed^5^ despite being granted full authorization in October 2020^6^. Monoclonal antibodies that target the viral spike protein have also been successful in treating disease when applied at the early stage of infection^7^. Though the availability of highly efficacious vaccines provides great hope for the eradication of COVID-19, it is expected that it will still take time for cases to subside globally and does not address the potential threat of future coronaviruses. Thus, other classes of antivirals that exhibit single agent efficacy or that are complementary to remdesivir for use in combination regimens are essential to meet this substantial unmet need.

SARS-CoV-2 produces two large viral polyproteins, pp1a and pp1ab, which are processed by two virally encoded cysteine proteases, the main protease, also called 3C-like protease (3CL protease or 3CLpro) and the papain-like protease. Mutagenesis experiments with other coronaviruses have demonstrated that the activity of the 3CLpro is essential for viral replication^8,9^. 3CLpro proteolytically processes the virus p1a/p1ab polyproteins at more than 10 junctions to generate a series of non-structural proteins critical for virus replication and transcription, including RdRp, the helicase, and the 3CLpro itself^10^. No close human analogs of the coronavirus 3CLpro are known, suggesting that selective 3CLpro inhibitors should avoid unwanted polypharmacology^11^. The essential functional importance of proteases in virus replication has led to the clinical success of protease inhibitors in combating both human immunodeficiency virus (HIV) and hepatitis C virus (HCV)^12–14^. This together with the opportunity for selectivity, makes 3CLpro an attractive antiviral drug target^15^.

Following the severe acute respiratory syndrome (SARS) outbreak in 2002-2003 we identified a potential small molecule protease inhibitor (PF-00835231) for the treatment of SARS-CoV-1, using structure-based drug design^16^. Due to the SARS pandemic being brought under control in July 2003 following public health measures which incorporated patient isolation and travel restrictions, this project was discontinued due to the lack of a path forward to demonstrate clinical efficacy. Given that the SARS-CoV-1 and SARS-CoV-2 3CLpro sequences share 96% identity overall and 100% identity in the active site^1,17,18^, following the emergence of SARS-CoV-2, PF-00835231 was identified as a potential SARS-CoV-2 3CLpro inhibitor for the treatment of COVID-19 disease^16^. Subsequent antiviral data presented here indicate that PF-00835231 has similar potency against either SARS-CoV-1 or SARS-CoV-2 (Table S1).

Herein we describe the 3CLpro inhibitor, PF-00835231, and its novel phosphate prodrug, PF-07304814, and present broad-spectrum inhibitory activity across coronaviral 3CL proteases, in vitro and in vivo antiviral activity in a coronavirus animal model, and absorption, distribution, metabolism, excretion (ADME) and safety data highlighting its potential for the intravenous (IV) treatment of COVID-19 disease.

## Results and Discussion

### PF-00835231 exhibits tight and specific binding to SARS-CoV-2 3CL *in vitro*

A thermal-shift assay was used to evaluate the direct binding between PF-00835231 and SARS-CoV-2 3CLpro. The melting temperature of SARS-CoV-2 3CLpro was shifted by 14.6℃ upon binding of PF-00835231, from 55.9±0.11℃ (n=16) to 70.5±0.12℃ (n=8). These data support tight and specific binding of PF-00835231 to SARS-CoV-2 3CLpro (Fig. 1) as was shown previously by X-ray co-crystal structure and evaluation of Ki ^16^.

**Fig. 1.**
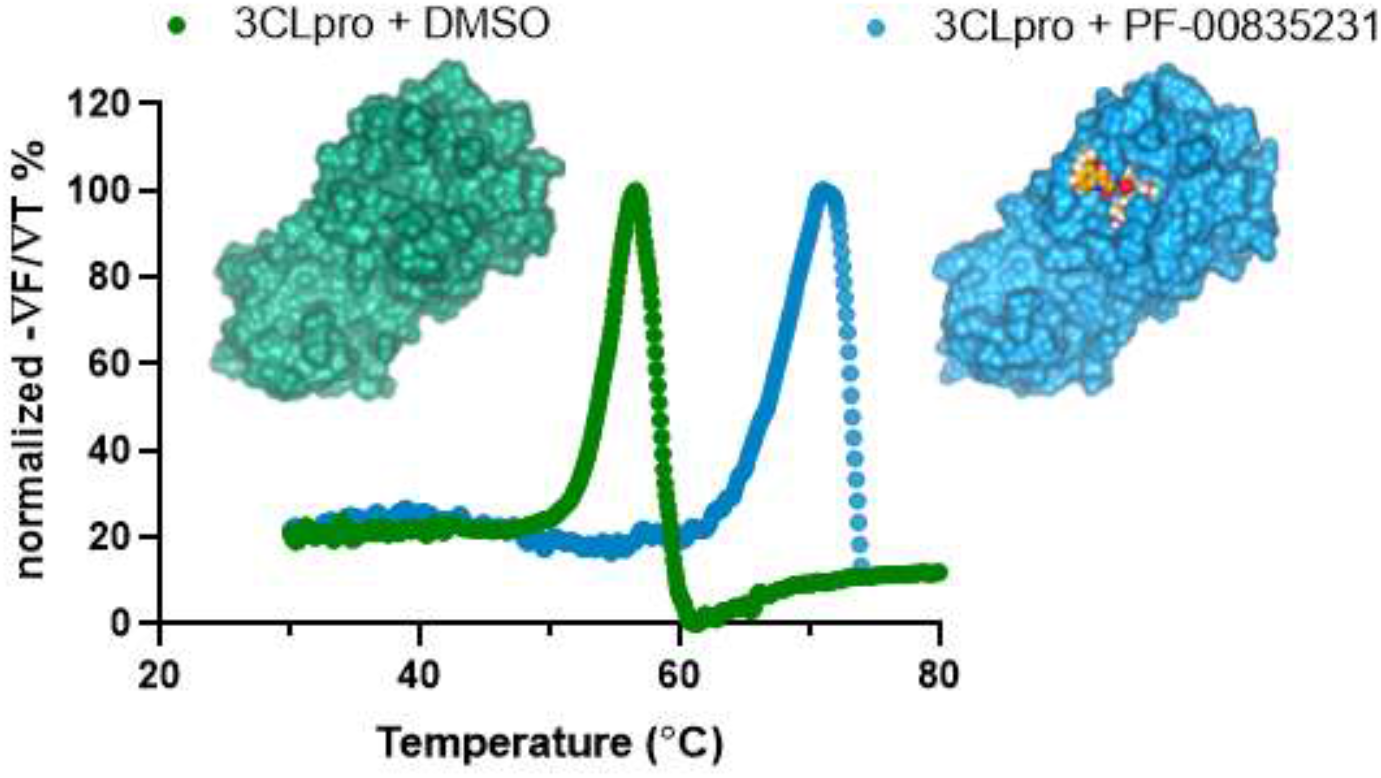
Representative thermal shift binding data of PF-00835231 with SARS-CoV-2 3CLpro. X-ray structures of SARS CoV-2 3CLpro apoenzyme (left) and SARS CoV-2 3CLpro in complex with PF-00835231 (right).

### PF-00835231 has potent and broad-spectrum inhibitory activity against a panel of coronavirus 3CLpros

To explore the notion that PF‑00835231 could have pan-coronavirus activity, PF-00835231 was evaluated against 3CLpro from a variety of other coronaviruses representing alpha, beta and gamma groups of *Coronaviridae*, using biochemical Förster Resonance Energy Transfer (FRET) protease activity assays. PF-00835231 demonstrated potent inhibitory activity against all tested coronavirus 3CLpro including members of alpha-coronaviruses (NL63-CoV, PEDV, FIPV), beta-coronaviruses (HKU4-CoV, HKU5-CoV, HKU9-CoV, MHV-CoV, OC43-CoV, HKU1-CoV), and gamma-coronavirus (IBV-CoV), with Ki values, ranging from 30 pM to 4 nM (Table 1). The demonstrated activity is consistent with a potential therapeutic use against emerging coronaviruses. This inhibitory activity is restricted to coronavirus 3CLpros as PF-00835231 was inactive against a panel of human proteases and HIV protease (Table S2). PF-00835231 showed detectable activity against human cathepsin B but 1000-fold weaker (6.9 nM vs 6 μM) activity compared to 3CLpro (Table S2). Thereby, these data collectively support PF-00835231 is a selective *in vitro* protease inhibitor with broad coronavirus activity.

**Table 1.** Activity of PF-00835231 against 3CLpro of coronaviruses

| **Virus** | **K<sub>i</sub> (nM)** |
| --- | --- |
| **Alpha-CoV** |  |
| NL63-CoV | 0.77 ± 0.52 |
| HCoV-229E | 1.5 ± 0.76 |
| PEDV | 0.30 ± 0.11 |
| FIPV | 0.12 ± 0.10 |
| **Beta-CoV** |  |
| SARS-CoV-2* | 0.27 ± 0.1 |
| HKU1-CoV | 0.85 ± 0.24 |
| HKU4-CoV | 0.034 ± 0.079 |
| HKU5-CoV | 0.033 ± 0.12 |
| HKU9-CoV | 0.74 ± 0.68 |
| MHV-CoV | 1.2 ± 0.90 |
| OC43-CoV | 0.51 ± 0.12 |
| **Gamma-CoV** |  |
| IBV-CoV | 4.0 ± 0.37 |

\*Data reported in <sup>16</sup>

### *In vitro* cellular antiviral activity of PF-00835231 against SARS-CoV-2

The antiviral activity of PF-00835231 against SARS-CoV-2 in cell culture was evaluated with a cytopathic effect (CPE) assay using either VeroE6 kidney cells enriched for angiotensin-converting enzyme 2 (ACE2) (VeroE6-enACE2) receptor or VeroE6 cells constitutively expressing EGFP (VeroE6-EGFP). These cell lines were infected with the SARS-CoV-2 Washington strain 1 (WA1 - EPI_ISL_404895) or the Belgium/GHB-03021/2020 strain (GHB-03021 - EPI_ISL_407976)^19^, respectively, which have identical 3CLpro amino acid sequences. PF-00835231 exhibited viral CPE EC_50_ values of 39.7μM and 88.9μM, respectively (EC_50_, Fig. 2). However, Vero cells express high levels of the efflux transporter P-glycoprotein (P-gp) (also known as MDR1 or ABCB1), of which PF-00835231 is a known substrate^16^ suggesting that the intracellular concentration of PF-00835231 was lower than it initially appeared. Therefore, to evaluate the full potency of PF-00835231, the assays were repeated in the presence of a P-gp efflux inhibitor, CP-100356^20^. PF-00835231 exhibited a 117- to 173-fold increase in activity in the presence of 2 μM P-gp inhibitor, with EC_50_ values of 0.23μM in VeroE6–enACE2 cells and 0.76μM in the VeroE6-EGFP cells (Fig. 2). The P-gp inhibitor alone had no antiviral or cytotoxic activity at these concentrations and did not cause cytotoxicity in the presence the protease inhibitor. The use of VeroE6 cells by many in the field to evaluate inhibitor activity could be problematic, since the true activity of some compounds could be masked by efflux of inhibitors from these cells (Fig. 2B). Consistent with many viral protease inhibitors^21^, there was a steep response to increasing doses of PF-00835231, with a ~2-3 fold difference between EC_50_ and EC_90_ in both cell types (EC_90_ = 0.48μM in VeroE6-enACE2 cells and EC_90_= 1.6μM in VeroE6-EGFP cells in the presence of the P-gp inhibitor). As expected, when lung cell lines were tested for antiviral potency in the presence and absence of P-gp inhibitor (A549-ACE2^22^ and MRC5), no statistical difference in antiviral potency was observed (Fig. 2A). Additionally, antiviral activities in both VeroE6 cell lines with 2μM P-gp inhibitor are similar to those observed in more physiologically relevant human lung cell culture systems, including A549-ACE2 and polarized human airway epithelial cells^22^, where P-gp expression is lower. These data support the potential for single agent antiviral activity. As will be presented later, despite seeing apparent antiviral activity for PF-07304814, a prodrug of PF-00835231, in cell-based assays (Fig. 2A) this activity is likely due to conversion of PF-07304814 to PF-00835231 in the assay.

**Fig. 2.**
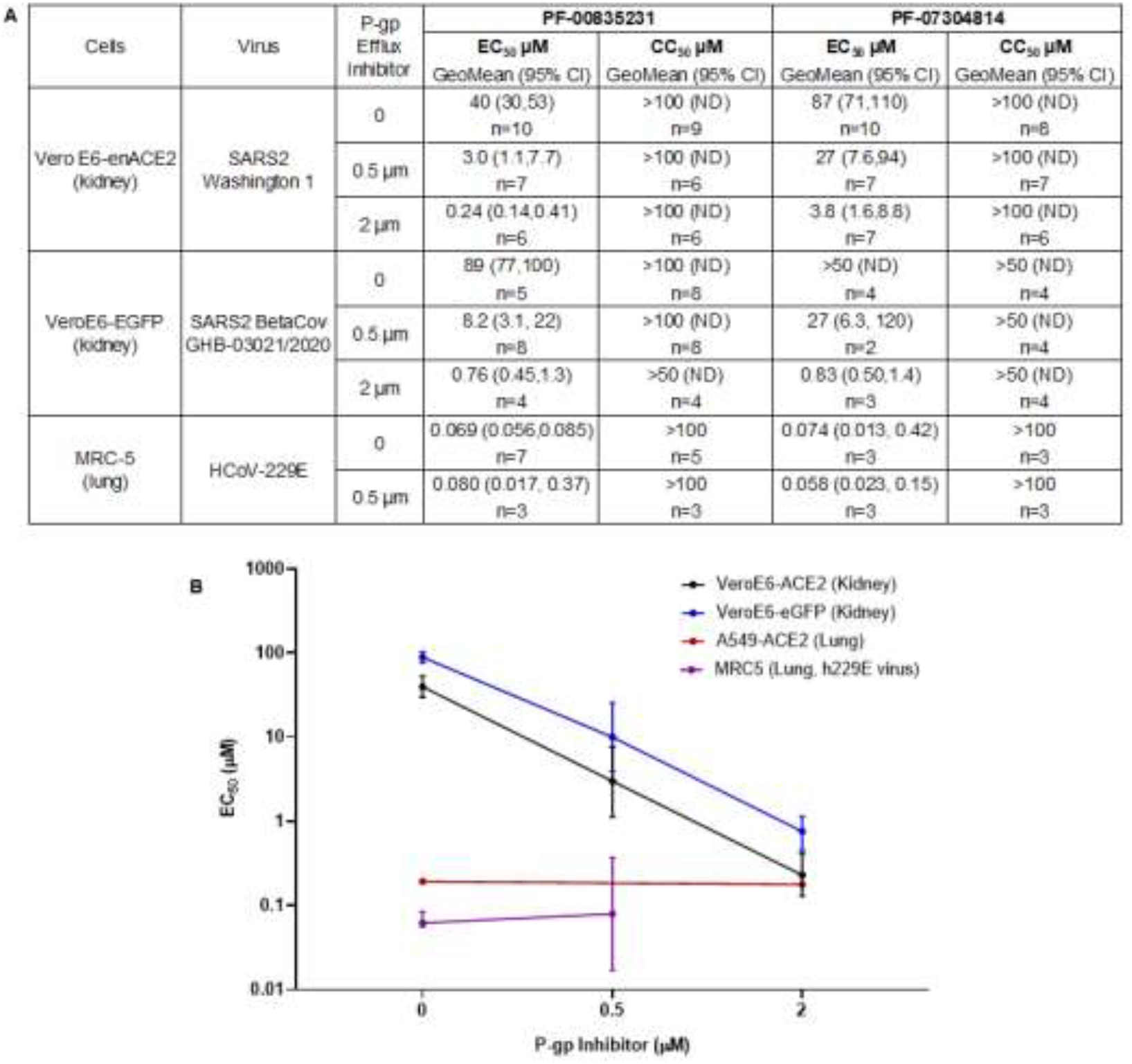
Antiviral activity across cell lines and viruses. (**A**) *In vitro* antiviral activity, and cytotoxicity for PF-00835231 and PF-07304814 with and without the P-gp efflux inhibitor, CP-100356. (**B**) EC_50_ values with PF-00835231 with increasing P-gp inhibitor in human lung and monkey kidney cell lines. A549-ACE2 human lung carcinoma data (red) as reported in ^22^.

### Potential for antiviral combination benefit of PF-00835231 in combination with remdesivir

Combinations of antiviral agents, especially those targeting different steps in the virus replication cycle, are a frequently employed therapeutic strategy in treating viral diseases^23^. As PF-00835231 and remdesivir, a nucleoside RNA-dependent RNA polymerase inhibitor, target different steps in the virus replication cycle, the antiviral activity of the two compounds was evaluated alone and in combination using HeLa-ACE2 cells^24^. Viral proteins were detected in this assay using convalescent human polyclonal sera from two different COVID-19 patients. PF-00835231 alone inhibited SARS-CoV-2 replication with an average EC_50_ of 0.13μM and EC_90_ of 0.43μM; whereas remdesivir had an average EC_50_ of 0.074μM and EC_90_ of 0.17μM (Fig. 3A). Combination studies were performed using a drug testing matrix, and the data for the drug combination were analyzed using reference models (Loewe, Bliss, HSA) to classify the effects of the drug combination as either additive, synergistic or antagonistic (isobologram, synergy scores, and combination indices).

**Fig. 3.**
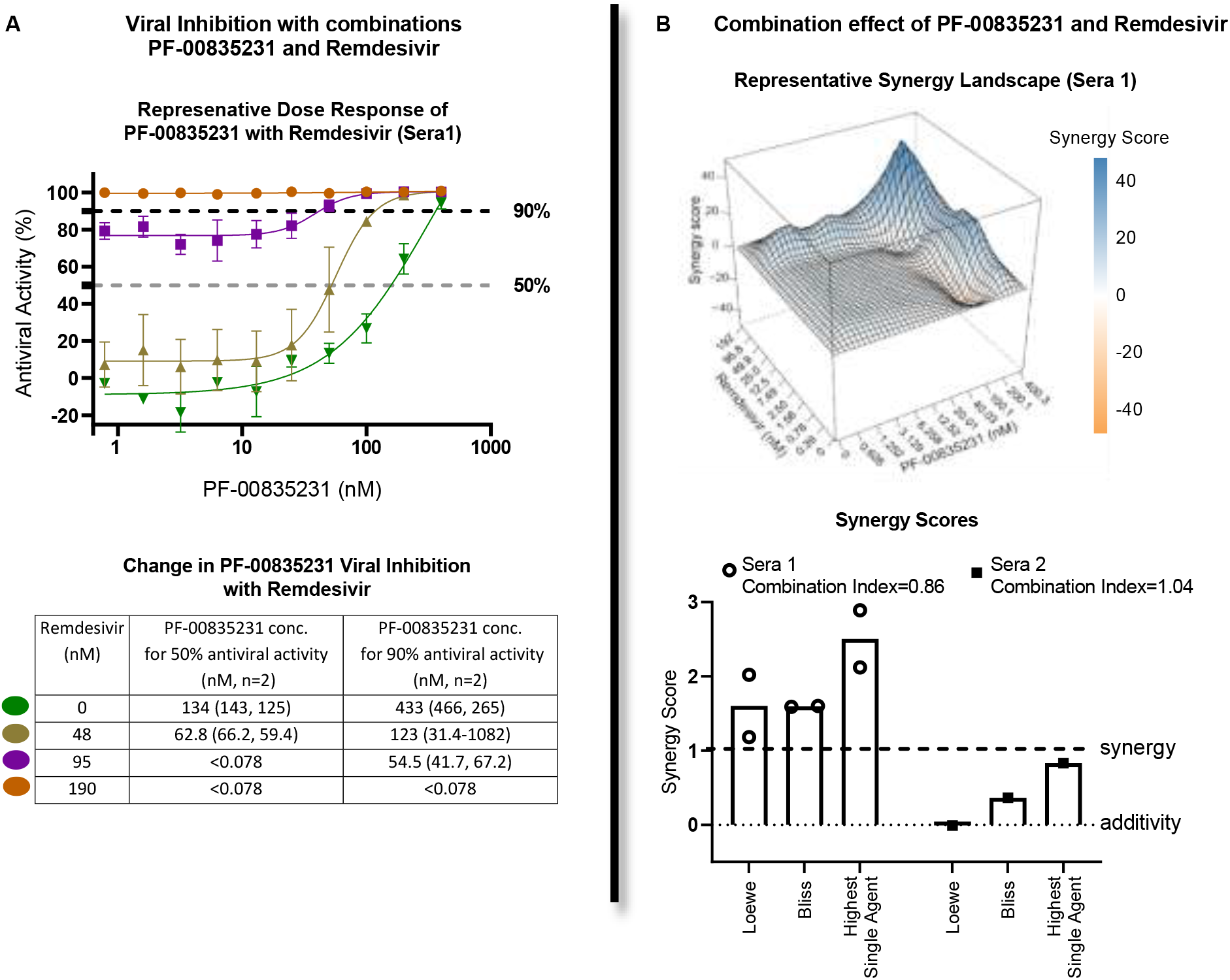
Measuring potential synergy between PF-00835231 and remdesivir in HeLa-ACE2 cells. (**A**) (Top) Representative antiviral dose response curves of PF-00835231 in combination with remdesivir against SARS-CoV-2. Serial dilutions of PF-00835231 with a range of fixed concentration of remdesivir. (Bottom) *In vitro* absolute antiviral activity shift in 50% and 90% antiviral activity with fixed concentrations of remdesivir. (**B**) (Top) A representative 3-dimensional drug interaction landscape plotting synergy scores analyzed using Synergyfinder (median scores of 3 replicates). (Bottom) Average *in vitro* combination synergy scores from the 3 experiments using 2 different patients’ sera (shown separately).

As summarized in Fig. 3B, the combination of PF-00835231 and remdesivir using patient sera for detection, exhibited synergy in two independent experiments with sera from patient #1 and additivity in a single experiment with sera from patient #2 (Fig. 3B). The different classification is most likely due to the different convalescent serum used as detection reagents. These same antiviral data were also analysed using Synergyfinder, which also indicated that the two drugs were additive to synergistic, with a representative graph shown in Fig. 3B. Antagonism was not demonstrated for the combination of PF-00835231 and remdesivir in these studies. The observed additivity/synergy was not due to cytotoxicity, as there was no noticeable cytotoxicity in virus infected host cells for all the combinations tested. This additivity/synergy is similar to other protease inhibitors used for the treatment of HCV, which has led to substantial clinical benefit^25^.

### Activity of PF-00835231 in a mouse model of SARS-CoV-1 infection

Human coronaviruses can replicate in mice but do not cause disease. The poor replication is due to differences between the human and mouse orthologs of the primary SARS-CoV-1 and CoV-2 receptor, ie, ACE2, unless the virus is first mouse-adapted (MA)^26^. Given the sequence identity between SARS-CoV-1 and SARS-CoV-2 3CL proteases, especially in the active site (100%), it was expedient to evaluate PF-00835231 in the validated SARS-CoV MA15 model^19^. SARS-CoV-MA15 has two mutations in 3CL protease coding sequence^27^ that are located distal to the active site that did not influence activity of the protease or its inhibition by PF-00835231 (Table S1). Treatment of MA15-infected BALB/c mice with 100 mg/kg PF-00835231 twice daily (BID) by subcutaneous (S.C.) injection, started either at the same time as infection, ie, day 0, or on day 1 or 2 post-infection (Fig. 4A). This resulted in a significant 1-2 log_10_ (Fig. 4B) reduction in lung viral titers, respectively, when measured on day 4. When the mice were treated with 30, 100, or 300 mg/kg (S.C., BID), a dose dependent reduction in lung viral titers of ≥3 log_10_ was observed (Fig. 4C). An assessment of plasma exposure was made following 100 mg/kg and 300mg/kg in non-infected animals and the unbound concentrations at 12h (Cmin) were approximately 500nM (~1x in vitro EC_90_) and 1700nM (~3x in vitro EC_90_) respectively. No assessment of the 30mg/kg exposures were made at the time of the experiment. Infected-mice that were treated with 300 mg/kg PF-00835231 starting on day 0 did not lose as much weight as vehicle-treated or infected, untreated animals (Fig. 4D). Microscopic evaluation of lung tissue showed that PF-00835231-treatment prevented or decreased histopathologic signs compared to vehicle-treated animals (Fig. 4E). Therefore, PF-00835231 was active *in vivo* when added at the time of infection or as treatment (delayed to day 1 or day 2, post-infection), alleviated signs of disease (weight loss and lung pathology), and drug exposure approximately equivalent to the in vitro EC_90_ was sufficient to achieve the maximal 2-3 log_10_ reduction in lung viral titers.

**Fig. 4.**
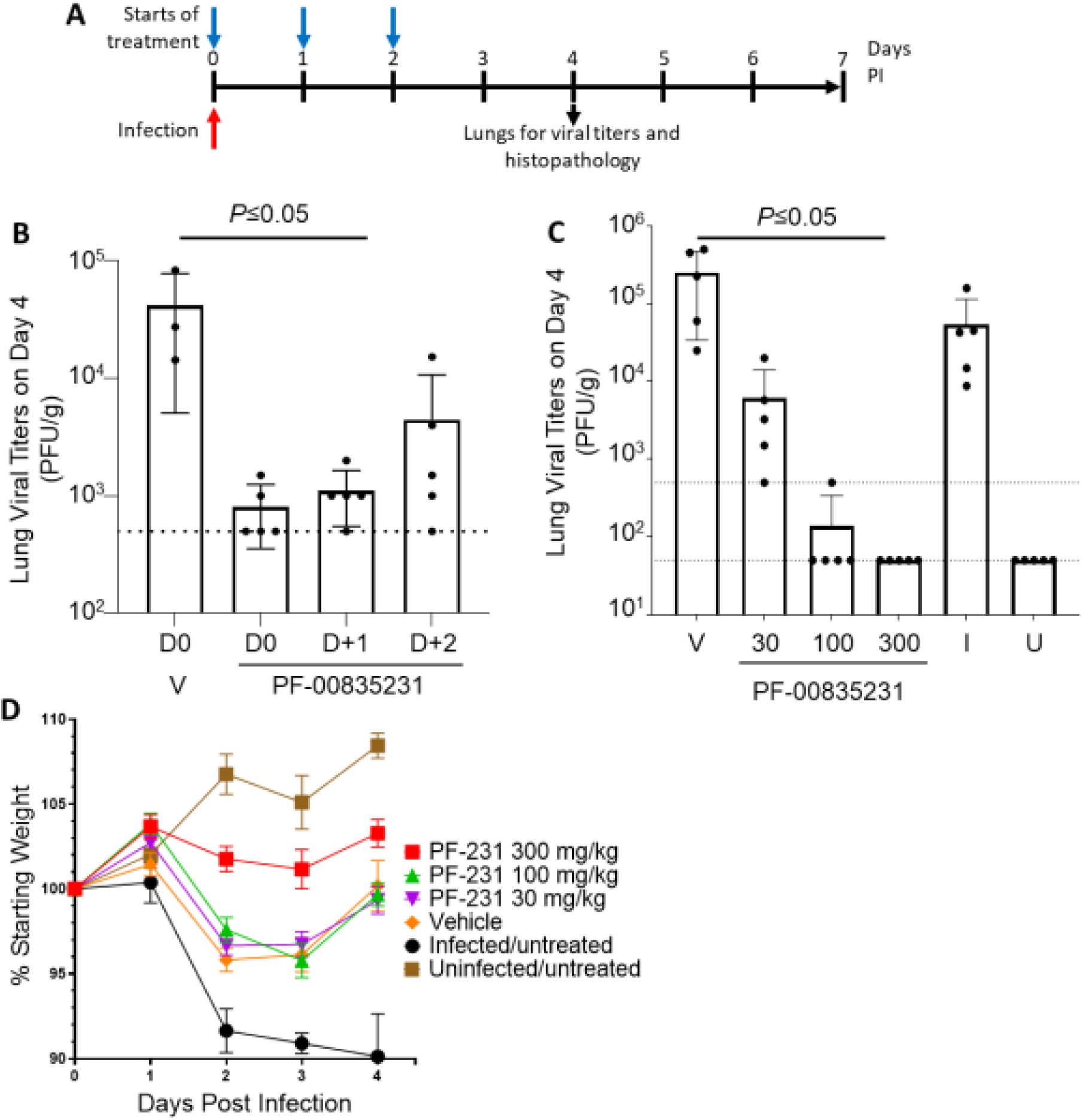
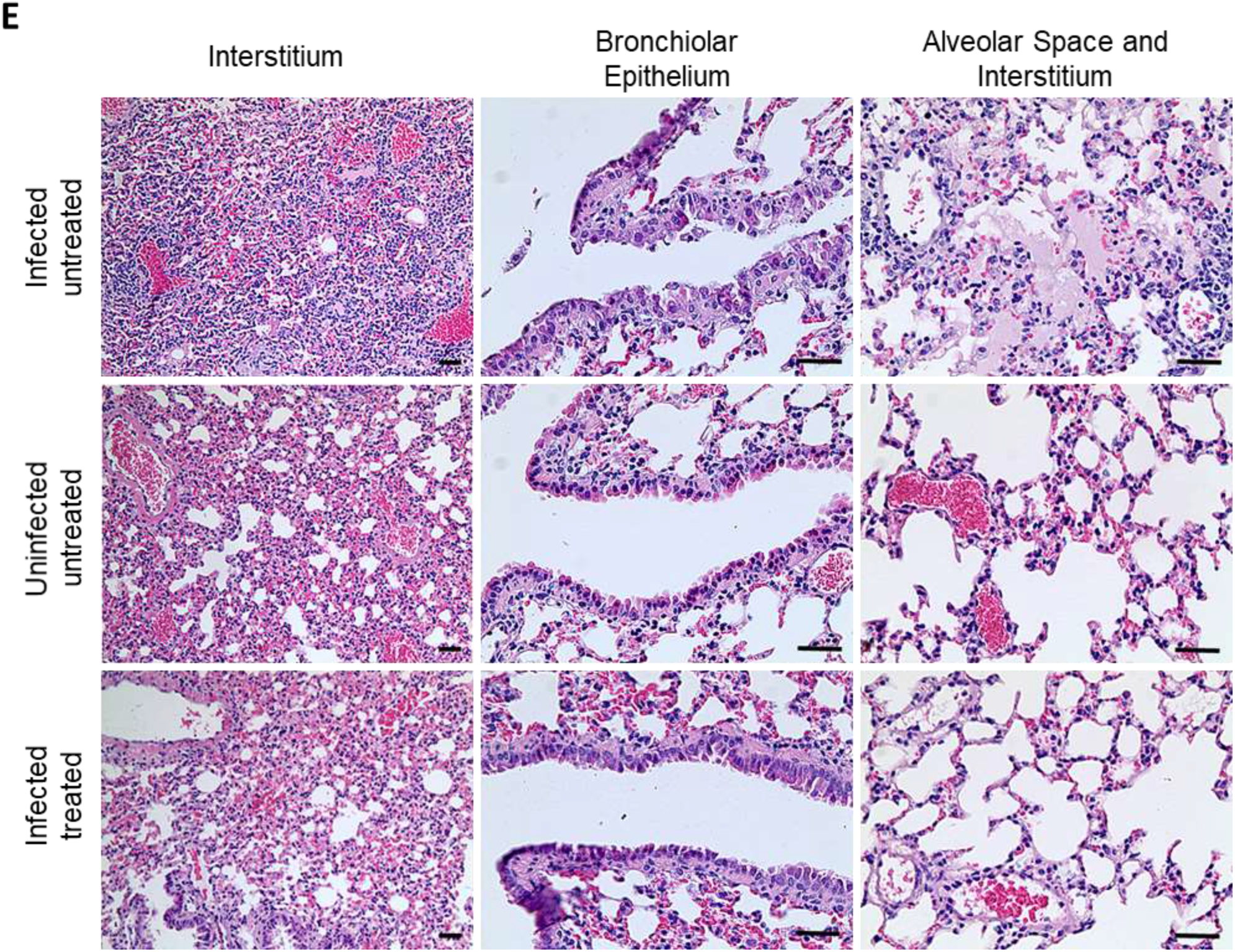
In vivo activity of PF-00835231 in a mouse model of SARS-CoV infection. **A**. Study design for in vivo experiments. Infection with SARS-CoV-MA15 was always on day 0. Treatment began on day 0 (panel C) or day 0, day 1, or day 2 post-infection (panel B). Lungs were harvested on day 4 post-infection for viral titers and lung histopathology (n=5). **B**. Lung viral titers for PF-00835231-treated (100 mg/kg, S.C. BID) mice when treatment was started on day 0 or delayed to day 1 and day 2 post-infection. **C**. Lung viral titers for mice treated in a dose-response of PF-00835231 at 30, 100, 300 mg/kg, S.C. BID starting on day 0. **D**. Change in body weight of the mice from the experiment shown in panel C. **E**. Representative photomicrographs of lung sections stained with hematoxylin and eosin. Lungs from infected/untreated mice (top row) displayed perivascular and interstitial inflammation (top, left), degeneration and desquamation of the bronchiolar epithelium (top, middle) and proteinaceous exudate in the alveolar space with interstitial inflammatory cells (top, right), all of which were not observed in uninfected/untreated mice (middle row), and in the infected mice treated with PF-00835231 at 300 mg/kg (bottom row). The scale bars represent 50 μm. LOD, limit of detection; I, infected, untreated; LOQ, limit of quantification; U, uninfected; V, vehicle-treated; Viral Titers error bars represent standard deviation; Weight change error bars represent SEM.

### Favorable preclinical ADME and pharmacokinetic profile of PF-00835231

The metabolic stability of PF-00835231 was evaluated *in vitro* using pooled human liver microsomes (HLM). PF-00835231 was shown to be metabolized by cytochrome P450 enzymes exhibiting an unbound CL_int_ 14μL/min/mg. Using recombinant heterologously expressed enzymes and HLM with the CYP3A selective inhibitor ketoconazole, CYP3A4 was identified as the major CYP involved in the metabolism of PF-00835231 (Table S3). It was also noted that the polymorphically expressed CYP3A5 can also metabolize PF-00835231 and that clearance may be slightly greater in CYP3A5 expressers. The potential for PF-00835231 to reversibly inhibit human cytochrome P450 enzymes (CYP1A2, 2B6, 2C8, 2C9, 2C19, 2D6, and 3A) was evaluated using probe substrates in pooled HLM and provided IC_50_ values >200 μM (Table S4) and a weak signal for time dependent inhibition (TDI) of CYP3A4/5 (Table S5). These data indicate PF-00835231 provides a low risk of causing drug-drug interactions (DDIs) on coadministration with other drugs. The potential for PF-00835231 to inhibit a range of transporters (BCRP, P-gp, OATP1B1/1B3, OCT1/2, OAT1/3 and MATE1/2K) was evaluated using *in vitro* systems (Table S6). The IC_50_ values were >20μM and indicate a low risk of causing DDIs due to transporter inhibition at the projected clinical exposure. The plasma protein binding of PF-00835231 was measured across species using equilibrium dialysis and demonstrated moderate binding to plasma proteins with plasma unbound fractions of 0.33 to 0.45 across species (Table S7).

PF-00835231 was administered IV to rats, dogs and monkeys (1 or 2mg/kg) and exhibited moderate plasma clearances (50-80% liver blood flow), low volumes of distribution (<1.5 L/kg) and short half-lives (<2 hours) across species in keeping with its lipophilic (LogD_7.4_=1.7) and neutral physiochemistry. Following oral administration to rats (2 mg/kg) and monkeys (5 mg/kg) PF-00835231 exhibited low oral bioavailability (<2%), likely due to a combination of low absorption because of its low permeability (apparent MDCK-LE permeability of 1.3×10^−6^cm/sec^28,29^), low solubility, potential for active efflux in the gut by P-gp and BCRP, and the potential for amide hydrolysis by digestive enzymes in the gastrointestinal tract. In rat, dog and monkey approximately 10% or less of PF-00835231 was eliminated unchanged in the urine indicating renal elimination may also play a minor role in the clearance of PF-00835231 in humans (Table S8).

### Human pharmacokinetic predictions suitable for IV administration

Taking into account the human *in vitro* metabolism data and *in vivo* pharmacokinetic (PK) data in rats, dogs and monkeys, PF-00835231 was predicted to exhibit a plasma clearance (CL_p_) of ~6mL/min/kg (major CYP, minor renal pathways), steady state volume of distribution (V_dss_) of 1L/kg and half-life of approximately 2 hours in humans. Due to the limited oral bioavailability, short elimination half-life, and the likely need to maintain unbound systemic concentrations over time to achieve antiviral activity, a continuous IV infusion was proposed as the optimal dosing route and regimen.

### Efficacious target concentration and feasible human dose projection

The projected minimally efficacious concentration (C_eff_) was chosen to match the *in vitro* EC_90_ (See supplemental methods for rationale), consistent with the preclinical to clinical translation of approved protease inhibitors^30^. Since PF-00835231 was proposed to be administered by continuous infusion, the projected steady-state exposure is equal to the C_min_ maintained over the dosing interval. The dose response assay performed in the most physiologically relevant cell type, human lung carcinoma, resulted in an average EC_90_ value of 0.44μM^22^. This is consistent with additional antiviral data in Hela-ACE2 cells (EC_90_=0.4μM) and Vero cell lines (EC_90_= 0.48-1.6μM) when a P-gp inhibitor was added to better reflect the lack of substantial P-gp transporter in the lung (Fig. 2B). Furthermore, the antiviral inhibition is supported by the antiviral time course experiment performed in a primary human airway epithelial model (preliminary data indicates an unbound EC_90_ <0.5 μM)^22^, indicating a consistent intrinsic anti-SARS-CoV-2 activity of PF-00835231 across different cell types. Therefore, the proposed minimal C_eff_ is ~0.5μM unbound.

Due to the rapid blood perfusion through the lungs and the continuous steady state intravenous infusion regimen, the unbound plasma and unbound lung concentrations are assumed to be in equilibrium and, therefore, the unbound plasma concentration provides a reasonable surrogate for the concentration at the main site of action of the disease. Based on the human PK predictions, the minimally efficacious dose of PF-00835231 necessary to achieve this exposure is ~320 mg/day administered as an intravenous continuous infusion. The required duration of dosing for efficacy remains uncertain and will need to be evaluated in humans.

### Formulation and solubility profile of PF-00835231 to enable IV administration

PF-00835231 is a moderately lipophilic (LogD_7.4_ = 1.7), neutral molecule with no ionizable centers throughout the physiologically relevant pH range. Consequently, PF-00835231 exhibits a pH independent solubility with an intrinsic aqueous solubility of less than 0.1mg/mL and limited opportunities for solubility-enabling formulation approaches. Preliminary work using standard solubilizing excipients indicated that achieving a solubility >0.5mg/mL would likely be challenging.

Based on a maximum desired intravenous infusion volume of ~1L per day a solubility of 0.5mg/mL would be sufficient to deliver the minimal efficacious dose estimate of ~320 mg/day to maintain a ~0.5μM steady state unbound concentration (Fig. 5B). Due to the nascent understanding of the virus, the required target levels of inhibition for clinical benefit remain uncertain and the ability to evaluate exposures up to ~10x C_eff_ in early clinical development is desirable. As a potential option to increase exposures, and/or decrease the required infusion volume, the use of a strong CYP3A inhibitor (itraconazole 200mg QD for 15 days) was considered but preliminary, physiologically-based pharmacokinetic (PBPK) modeling predicted only a ~2-fold increase in PF-00835231 exposure at steady state (Table S11).

**Fig. 5.**
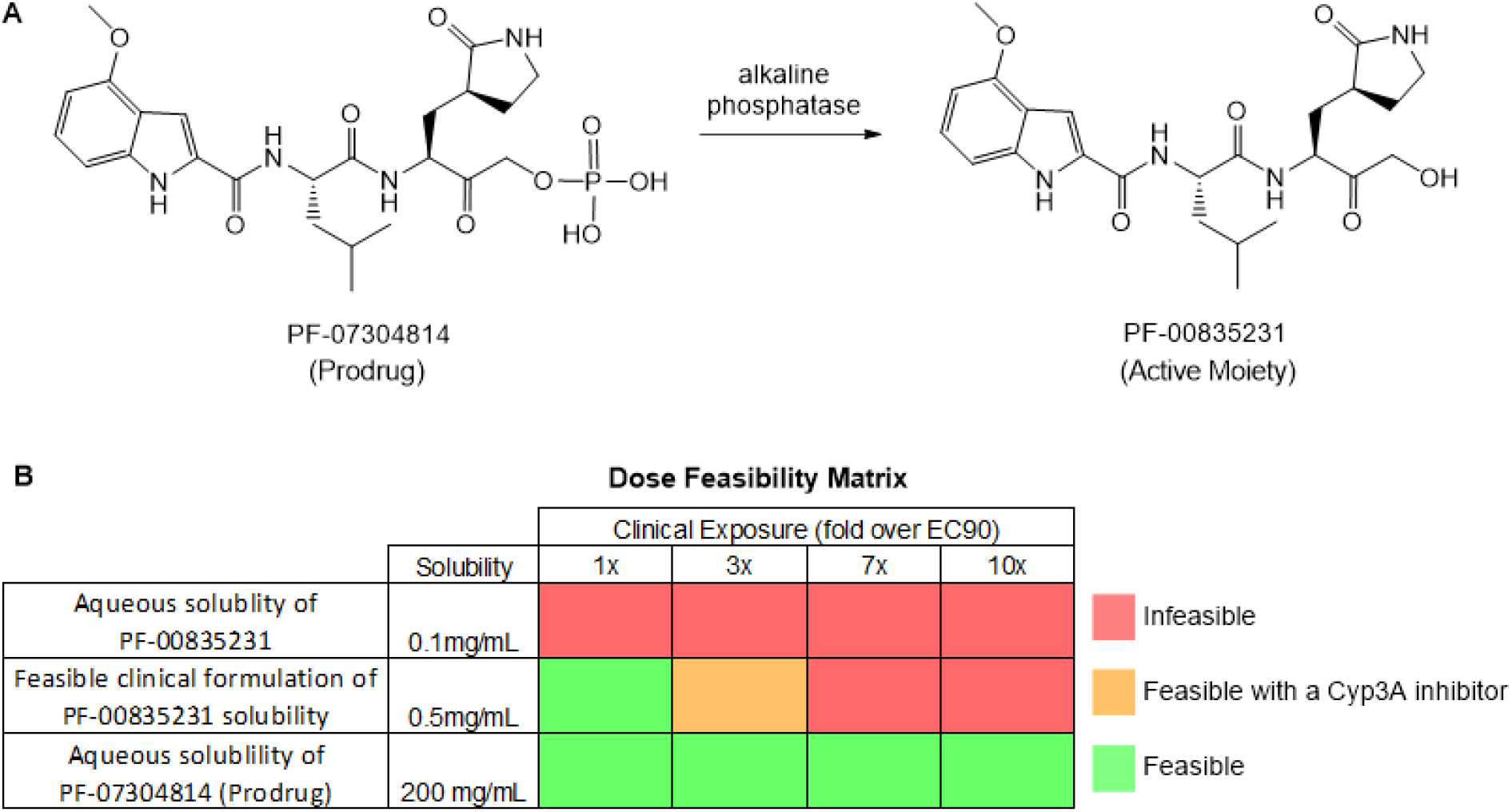
PF-7304814 prodrug and PF-00835231 structures and dose considerations. (**A**) Chemical structure of conversion of prodrug PF-07304814 to the active moiety PF-00835231 by alkaline phosphatase. (**B)** Dose feasibility matrix illustrating the ability to achieve higher target exposures with increasing solubility and the limitations of dosing PF-00835231. with dosing either aqueous PF-00835231, clinically formulated PF-00835231, or aqueous PF-07304814 (prodrug). The infeasible limit (red) is assumed to be 1 L per day with a 2x potential benefit with a Cyp inhibitor (orange). Any dose under that is considered feasible (green).

The ability to achieve higher doses could also potentially mitigate a higher than predicted clearance, or variations in patient body weight. Therefore, a medicinal chemistry strategy to significantly enhance the aqueous solubility of PF-00835231, by designing a phosphate prodrug was pursued.

### Considering an intravenous phosphate prodrug approach to improve solubility

IV phosphate prodrugs have precedence with several commercially available drugs such as fosfluconazole and fosphenytoin which are rapidly cleaved by human alkaline phosphatase to provide high systemic exposures of their respective active moieties following IV administration (Fig. 5) ^31,32^. Alkaline phosphatase is ubiquitously expressed across tissues with high levels expressed in the liver, lung, and kidney (Alkaline phosphatase tissue data from v19.proteinatlas.org^33^). High levels of conversion from prodrug to active moiety for fosfluconazole and fosphenytoin have also been observed in rats and dogs supporting cross species translation to human for the conversion of prodrug to active moiety^32,34^. Overall, the use of a phosphate prodrug is an established approach for IV administration to provide rapid conversion to its active moiety and was considered for PF-00835231.

### Synthetic route to provide phosphate prodrug

The synthesis of PF-00835231 has been described previously^16^. The subsequent synthesis of the phosphate prodrug of PF-00835231 was achieved via two steps (Fig. S1). Briefly, treatment of **1** (PF-00835231) with di-*tert*-butyl *N*,*N*-dipropan-2-ylphosphoramidite and tetrazole in tetrahydrofuran followed by oxidation with aqueous hydrogen peroxide delivered intermediate **2**. The phosphate *t*-butyl groups were subsequently hydrolyzed using trifluoroacetic acid in dichloromethane to deliver phosphate prodrug **3** (PF-07304814) as a solid.

### Enhanced formulation and solubility profile of PF-07304814 to provide clinical flexibility

PF-07304814 rapidly undergoes *in vivo* conversion to the active metabolite PF-00835231 (Fig. S2A). The phosphate prodrug is weakly acidic, with pKas of 1 and 6.4, and a predicted LogD_7.4_ of −3.7. At pHs above the compound’s first pKa, the phosphate functional group is de-protonated and negatively charged, which enables a significant improvement in aqueous solubility to greater than 200mg/mL over a pH range compatible with intravenous infusion. The higher intrinsic solubility of PF-07304814 eliminates the need for solubility-enabling formulations and enables the use of standard IV compatible excipients. Furthermore, the improved solubility enables higher doses to be explored in the clinic and gives clinicians greater flexibility in terms of dose volume to account for patient-specific co-administration requirements.

### PF-07304814 (prodrug) preclinical *in vitro* and *in vivo* ADME profile

To understand the metabolic stability and conversion of PF-07304814 to its active moiety (PF-00835231), PF-07304814 enzyme kinetics were evaluated *in vitro* using liver S9 fractions and was shown to exhibit rapid conversion to PF-00835231 with unbound CL_int_ values of 51, 84, 168 and 428μL/min/mg in rat, dog, monkey and human respectively. In these *in vitro* systems, PF-00835231 was the only metabolite formed from PF-07304814. Conversion was rapid in phosphate-free incubations but abolished in the presence of phosphate buffer supporting the role of alkaline phosphatase in this conversion (Table S9, Fig. S2). To evaluate the *in vivo* conversion and systemic availability of the active moiety PF-00835231, PF-07304814 was administered intravenously to rats, dogs and monkeys. PF-07304814 exhibited high conversion to PF-00835231, systemic clearance and short half-life across species forming 68, 81, 76% PF-00835231 in rats, dogs and monkey, respectively, in comparison to the systemic exposure achieved with IV administration of PF-00835231 (Fig. 6A and Table S8).

**Fig. 6.**
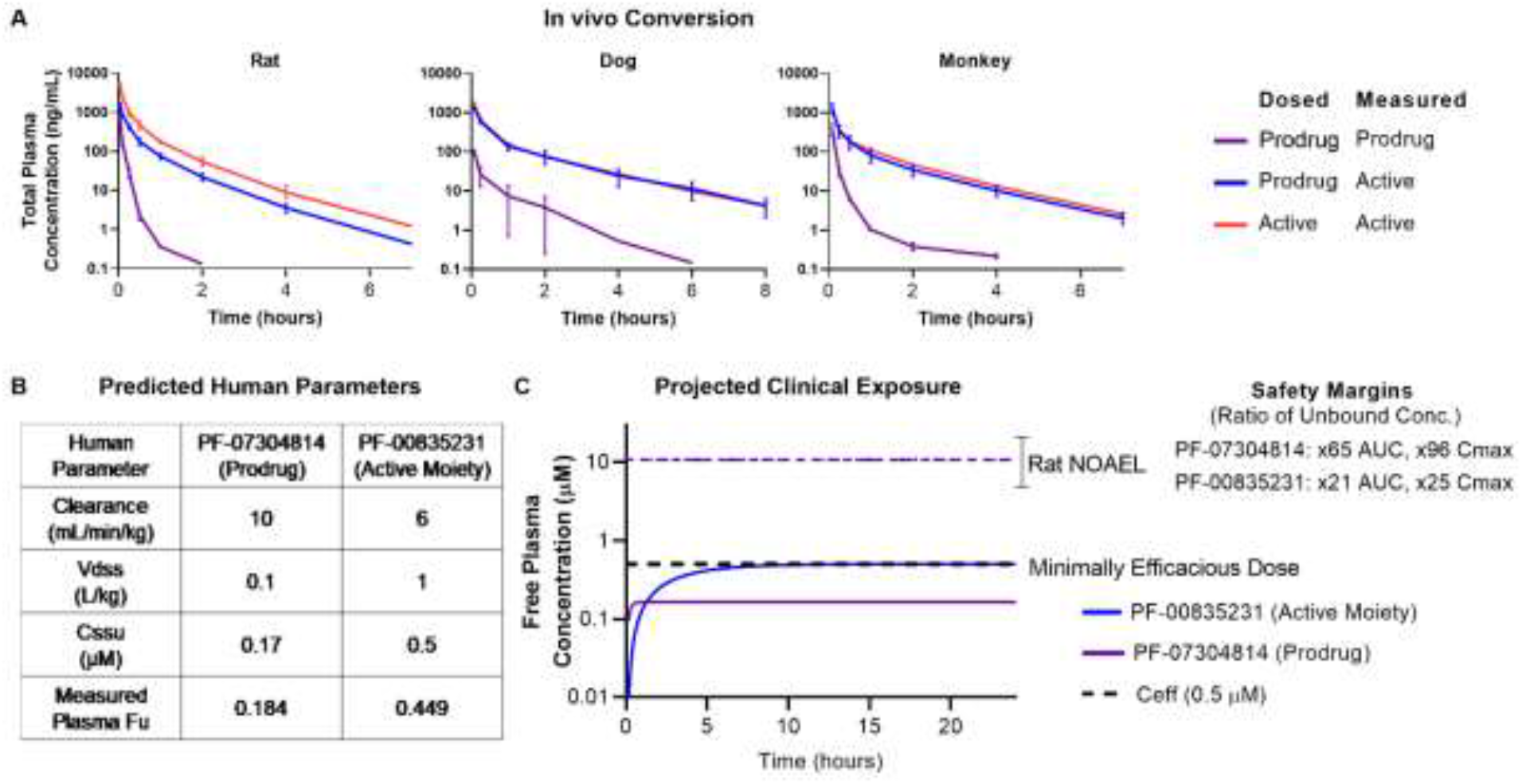
PF-07304814 (prodrug) and PF-00835231 *in vivo* exposure summary. (**A**) Rat, dog and monkey PK following IV administration of PF-07304814 (1.17mg/kg) or PF-00835231 (2mg/kg rat, 1mg/kg dog and monkey) demonstrating high levels of PF-00835231 formed *in vivo*. (n=2) (**B**) Predicted human PK parameters and measured protein binding for PF-07304814 and PF-00835231 used for human dose prediction. (**C**) Projected human systemic exposure profiles at the minimally efficacious dose of 500mg/day of PF-07304814 delivered as a continuous IV infusion. The predicted unbound steady state concentrations for the prodrug PF-07304814 (purple) and the active moiety PF-00835231 (blue) are 0.17μM and 0.5μM respectively. (NOAEL= No Observed Adverse Effect Level; C_eff_= projected minimally efficacious concentration, Fu= unbound fraction; error bars= range of replicates)

PF-07304814 was also evaluated for the potential to cause reversible and time dependent inhibition of human cytochrome P450 enzymes using pooled HLM and probe substrates for a range of CYP enzymes (CYP1A2, 2C8, 2C9, 2C19, 2D6, and 3A4/5) and showed low risk with IC_50_ values >100μM and no evidence of TDI (Table S4). The potential for PF-07304814 to inhibit a range of transporters (BCRP, P-gp, OATP1B1/1B3, OCT1/2, OAT1/3 and MATE1/2K) was evaluated using *in vitro* systems providing IC_50_ values >130μM indicating a low risk of causing DDIs due to transporter inhibition at the projected C_eff_ (Table S6) The plasma protein binding of PF-07304814 was measured across species using equilibrium dialysis showing moderate binding to plasma proteins with plasma free fractions of 0.18 to 0.38 across species (Table S7).

### Encouraging human PK predictions for PF-00835231 formation

The predicted human plasma clearance of PF-07304814 is ~10mL/min/kg based on scaling *in vitro* human liver S9 CL_int_ data (using equations 9 and 10, see Methods) and represents a conservative prediction of total Cl_p_ as it only accounts for conversion of prodrug to active moiety in the liver. The human Vd_ss_ for PF-07304814 is predicted to be ~0.1L/kg based on its acidic physiochemistry and observed human Vd_ss_ values of other phosphate prodrugs^31,32^. Based on the predicted Cl_p_, Vd_ss_, and a ~75% conversion to PF-00835231 based on the mean conversion in animals, PF-07304814 is anticipated to exhibit a short half-life of ~0.1hour with high conversion to the active moiety (Fig. 6B).

### PF-07304814 unlikely to contribute to antiviral activity *in vivo*

In a direct comparison, using the same SARS-Cov-2 3CLpro assay method as described in^16^, the prodrug PF-07304814 binds and inhibits SARS-CoV-2 3CLpro activity with a Ki of 174nM providing a >600-fold less potent Ki in comparison to the active moiety PF-00835231 (0.27nM)^16^. However, PF-07304814 shows apparently similar antiviral activity to PF-00835231 (1-12-fold, Fig. 2A) across cellular *in vitro* assays. This is most likely due to the partial conversion of PF-07304814 to PF-00835231 in the cellular assays by alkaline phosphatase. This was consistent with PF-00835231 concentrations measured at approximately 50% of the PF-07304814 starting concentration at the end of the 3-day incubation in the VeroE6 cell assay. It is not believed that PF-07304814 has antiviral activity without first converting to PF-00835231.

### PF-07304814 dose projection provides clinical flexibility to achieve target C_eff_

The antiviral activity and projected exposure for PF-00835231 after conversion from PF-07304814 *in vivo* was used to derive the minimal C_eff_ and dose estimates. Based on the predicted human PK and 75% conversion of the prodrug, a free plasma concentration of the active moiety PF-00835231 of 0.5μM (C_eff_) can be achieved with a 500 mg continuous IV infusion of the prodrug over 24 hours (Fig. 6C). The estimated time to achieve 90% steady state exposure of PF-00835231 is approximately 6 hours. Due to the improved solubility (>200mg/mL), the dose of PF-0730814 can be delivered in a volume of less than 0.25L. In addition the dose can be increased if the observed human plasma Cl exceeds 6mL/min/kg, if the percent converted from prodrug to active is less than predicted, or if exposures in excess of the minimal C_eff_ (0.5μM free) are required to maximize clinical activity (Fig. 5B). Overall, the improved solubility of PF-07304814 would theoretically enable >100-fold the proposed minimal C_eff_ dose in a 0.25L dose volume.

### Preclinical safety profile supports progression to clinical evaluation

A toxicology assessment consisting of an *in vitro* battery of genetic toxicity, secondary and safety pharmacology studies, in conjunction with a single species (rat) *in vivo* Good Laboratory Practice (GLP) study has been completed.

The safety profiles of PF-07304814 and PF-00835231 were assessed individually in a range of *in vitro* and *in vivo* safety studies in rats. In the *in vitro* studies, PF-07304814 and PF-00835231 were negative in the bacterial reverse mutation assay and did not induce micronuclei formation. Both compounds had minimal potential for secondary (off-target) pharmacology at clinically relevant exposures. Neither PF-07304814 nor PF-00835231 inhibited hERG current amplitude at up to 300μM (1,770- and 600-fold, respectively, in reference to the projected unbound human C_max_ of 0.17 and 0.50μM, respectively, at the projected human efficacious dose), indicating a favorable cardiovascular safety profile. In human blood hemocompatibility assays, both compounds had no effect on hemolysis or flocculation/turbidity parameters, indicating compatibility with human blood and supporting intravenous administration.

PF-07304814 was administered to rats via continuous IV infusion for 24 hours in a GLP study. There was no test article related findings and no target organ toxicity was identified. PF-07304814 had no effects on neurological safety pharmacology parameters as assessed by functional observation battery in the 24-hour continuous IV infusion rat study. The no observed adverse effect level (NOAEL) was 1000mg/kg. PF-00835231 was also administered to male rats via continuous IV infusion for 4 days in a non-GLP exploratory toxicity study and was tolerated at 246mg/kg/day, the highest feasible dose tested. PF-00835231-related findings in this study were limited to minimal, non-adverse effects on clinical chemistry parameters including higher mean triglycerides (1.9-3.6x vs controls), cholesterol (1.3x), and phosphorus (1.1x) without any microscopic correlates or associated functional changes. No test article related adverse effects were seen in any study.

At the NOAEL from the 24-hour, GLP continuous IV infusion study with PF-07304814 in rats, the anticipated exposure margins for unbound C_max_ and AUC_24_ are 97 and 65-fold for PF-07304814 and 25 and 21-fold for PF-00835231, at the projected minimum human efficacious dose of 500mg/day. This indicates the potential to safely evaluate multiples over EC_90_ in humans during clinical testing to understand the exposure response relationship and to achieve high levels of inhibition, if required. Furthermore, no overlapping or additive toxicity with medications currently being used in standard of care COVID-19 treatment is expected with administration of PF-07304814 in humans, making PF-07304814 an attractive combination partner. Based on results from the set of safety studies conducted, PF-07304814 exhibited an encouraging nonclinical safety profile and supported progression into Phase 1 clinical studies.

## Conclusions

PF-07304814 is a phosphate prodrug that is rapidly converted *in vivo* to the active moiety, PF-00835231, which exhibits high selectivity over human proteases, acts as a broad-spectrum coronavirus 3CL protease inhibitor, and demonstrates potent antiviral activity *in vivo*. Robust antiviral activity was demonstrated in a range of cellular *in vitro* assays in keeping with SARS-COV-2 human airway epithelial data^22^ suggesting a C_eff_ value of ~0.5μM unbound. This Ceff exposure was consistent with the free plasma concentration associated with the maximal viral load decrease in the in vivo mouse SARS-CoV model. The predicted human pharmacokinetics of PF-07304814 provide the ability to achieve systemic unbound concentrations of 0.5μM (EC_90_) of PF-00835231 by delivering 500mg as a continuous infusion over 24 hours with infusion volumes of less than 0.25L. In addition, higher doses (up to and beyond 10x C_eff_) also remain feasible due to the high solubility of PF-07304814.

Overall, PF-07304814 exhibits an encouraging preclinical profile that has the ADME, safety, and once converted to PF-00835231, SARS-CoV-2 antiviral activity to support progression to the clinic as a novel COVID-19 single agent antiviral treatment, with potential for further additional benefit in combination with antivirals that target other critical stages of the coronavirus life cycle. The favorable profile of PF-07304814 warrants clinical evaluation and is currently in clinical trials: NCT04627532 and NCT04535167.

## Acknowledgments

The authors would like to thank Sarah Lazzaro, Sumathy Mathialagan, Sangwoo Ryu, Mark West and Emi Yamaguchi (Pfizer) for the transporter inhibition studies. Angela Doran, Chad Limanni, Amanda Plante and Jocelyn Rosado for their in vivo and PK study support (Pfizer). Marcus Ewing (Pfizer) and Gail Johnson (Pfizer) for preformulation studies. Li Hao (Pfizer) for sequence analysis support. Shinji Yamazaki (Pfizer) for PBPK modeling simulations. Daniel Lettiere, Michael Homiski, Michelle Kenyon, Asser Bassyouni, Declan Flynn, William Reagan, Victoria Markiewicz and Stephen Jenkinson for overseeing safety studies, and William Reagan, for expert clinical pathology and pathology support for the toxicology studies. Deli Huang for supplying the HeLa-ACE2 stably transfected cell line.

## Author contributions

A.D.M, E.L and B.A contributed to conceptualization, investigation, analysis, visualization and data curation of inhibition of protease activity against a panel of coronavirus 3CLpro experiments. A.D.M contributed to funding acquisition, supervision, project administration and resources of these experiments. J.W and C.M contributed to conceptualization, investigation, analysis, visualization and data curation of the thermal shift experiments. JW contributed to funding acquisition, supervision, project administration and resources of these experiments. M.A.B, T.F.G supervised, designed and carried out antiviral synergy infection experiments. N.B, M.G.K and E.C contributed to the generation of *in vitro* antiviral synergy data J.B, J.H, Y.Z, L.M.A, L.L,S.N, R.O, C.S, R.K, R.H and B.B contributed to the analysis, interpretation of protease and antiviral data from collaborators and internal data. E.K designed and interpreted the ADME transporter experiments, R.S.O, H.E, R.M.J, E.P.K contributed to the metabolism, pharmacokinetics and bioanalysis of PF-07304814 and PF-00835231. J.R.L, S.A.L, M.N.O., and M.T. designed and interpreted formulation experiments and characterization of API properties. M.R.R, M.P, K.O, R.H and D.O designed and synthesized PF-07304814. R.M.J, B.B, R.S.O and E.K contributed to the conceptualization, analysis and calculations for the prediction of human PK and dose estimate for PF-07304814. J.G.S, L.W.U, R.M.J contributed to the design, supervision and interpretation of in vitro and in vivo safety study data. C.A, A.A, M.I.R contributed to the scientific discussion, experimental design, data interpretation in addition to manuscript review and editing. N.S. contributed to the analysis, interpretation of protease and antiviral data. S.W.M. and M.B.F. designed and interpreted and H.H., J.L., M.M., R.H., R.E.H., S.W. executed the in vivo animal efficacy studies. All authors contributed to writing drafts of the manuscript.

## Data and materials availability

All data are available in the main text or the supplementary materials.

## List of Supplementary Materials

Materials and Methods

Figures S1-S2

Tables S1-S11

References (35-65)

## Supplementary Materials

## Materials and Methods

### Thermal Stability Assay

3CLpro protein (3μM) was incubated with DMSO or 40 μM PF-00835231 in reaction buffer (20mM HEPES, pH 6.5, 120mM NaCl, 0.4mM EDTA, 4mM DTT and 20% glycerol) at 30 °C for 30 minutes. 1X SYPRO orange dye was added and fluorescence of the well was monitored under a temperature gradient range from 30°C to 90°C with 0.05 °C/s incremental step. The melting temperature (T_m_) was calculated as the mid-log of the transition phase from the native to the denatured protein using a Boltzmann model in Protein Thermal Shift Software v1.3.

### Coronavirus protease panel

#### Expression and purification of Alpha-, Beta-, and Gamma-coronavirus 3CLpro enzymes

In general, the coding regions for all 14 of the 3C-like proteases described in this study were codon-optimized and synthesized by a commercial source (BioBasic or GeneScript). The coding regions were inserted into a derivative of the pET 11 or pET15 expression vectors that place a hexa-histidine tag at the N-terminus that is followed by a Tobacco-Etch Virus (TEV) protease cleavage site that resides between the coding region of 3CLpro and the hexa-histidine affinity tag. The cleavage site for TEV is also recognized by 3CLpro, and therefore it is autocatalytically cleaved during expression in *Escherichia coli*, releasing authentic 3CLpro for purification. Each of the 14 coronaviruses were expressed in either *E. coli* BL21(DE3) or *E. coli* BL21(DE3) Gold cells and then purified after lysis using a combination of ammonium-sulfate fractionation, hydrophobic-interaction, anion-exchange, cation-exchange and size-exclusion chromatography. With the exception of 3CLpro from h229E, the details of the purifications for the other 13 3CLpros have been described in the literature with some modifications as outlined below.

The expression and purification of 3CLpro from SARS-CoV-2 3CLpro ^35^ SARS-CoV-1 ^36,37^, MERS ^38^, HKU4 ^39^, HKU5 ^40^, FIPV ^41^, PEDV ^42^, and MHV ^43^ have been previously described. Expression and purification protocols for 3CLpro from NL63, OC43, HKU9 and IBV coronaviruses were performed as described^44^ with the following modifications. NL63 3CLpro was expressed and purified using a modified protocol based on the purification of murine hepatitis virus (MHV) 3CLpro using a DEAE-Cellulose column. OC43 and HKU9 3CLpro were expressed using the autoinduction protocols developed for MERS 3CLpro followed by the same purification method as MHV 3CLpro. Avian coronavirus (IBV) 3CLpro was expressed and purified using a modified protocol based on MERS 3CLpro.

Expression and purification of the human h229E coronavirus 3CLpro was performed using the following method. The sequence for human h229E was obtained from UniProt (accession AGW80931.1) and was designed to include an N-terminal hexa-histidine tag followed by polyprotein residues 2960 - 3267. This gene sequence was codon-optimized for expression in *E. coli* and subcloned into a pET11a vector using the synthetic DNA services of GenScript Biotech (Piscataway, NJ). The pET11a plasmid containing the 229E 3CLpro gene was electroporated into *E. coli* BL21-GOLD (DE3) cells. A single colony of the transformed cells was used to inoculate 100mL of 2xYT media (10 g yeast extract, 16 g tryptone, 5 g NaCl per 1 L of water, 100 μg/mL carbenicillin, pH adjusted to 7.50 using 10 M NaOH). The preculture was incubated at 37°C until the culture reached an OD_600_ of 0.6. 15mL of the preculture was added to 1 L of 2xYT expression media and incubated at 37°C until an OD_600_ of 0.6 was reached. The cultures were then placed on ice for 15 minutes before addition of isopropyl B-D-1-thiogalactopyranoside (IPTG) to a final concentration of 0.6mM and further incubated at 25°C. After 16 hours, the cells were harvested via centrifugation at 10,940 x g for 10 minutes to yield an 8.3 g cell pellet per L. The cell pellet was resuspended in 5mL Lysis Buffer (25mM HEPES, 0.05mM EDTA, 5mM β-mercaptoethanol (β-ME), 1mg/mL lysozyme) per 1 g of pelleted cells using a manual homogenizer. The homogenized cell suspension was sonicated for a total of 12 minutes at an amplitude of 60% for periods of 10 seconds with 20 second delays using a Branson digital sonifier. Solid ammonium sulfate was slowly added to the resulting lysate to a final concentration of 1 M and stirred at 25°C for 10 minutes. The lysate was then clarified via centrifugation at 20,442 x g for 45 minutes at 4°C.

The resulting supernatant was checked for activity prior to being applied to a 75mL Phenyl Sepharose (GE Healthcare) resin equilibrated in Buffer B (1 M ammonium sulfate, 50mM HEPES, pH 7.50, 0.05mM EDTA, 5mM βME). The protein was eluted off the column using a linear gradient to 100% Buffer A (25mM HEPES, 0.05mM EDTA, 5mM β-ME) over 5 column volumes, followed by a water wash. 2mL fractions that eluted in water containing 229E 3CLpro were mixed with 8mL of 5X Buffer A. Fractions were assessed for purity via SDS-PAGE and enzymatic activity based on specific activity prior to pooling. The resulting protein pool was filtered using a 0.45μm surfactant-free cellulose acetate membrane prior to injection onto an 8mL MonoQ column equilibrated in Buffer A. The protein was eluted using a linear gradient to 100% Buffer C (1 M NaCl, 50mM HEPES, pH 7.50, 0.05mM EDTA, 5mM βME) over 15 column volumes. Fractions were pooled based on purity and specific activity and concentrated to 2.5mL using a 10,000 molecular weight cutoff (MWCO) spin concentrator (MilliporeSigma). The sample was then injected onto an SRT SEC-300 column (Sepax Technologies) that had been equilibrated in Buffer D (50mM HEPES, pH 7.50, 10% glycerol, 2.5mM dithiothreitol (DTT)). The resulting protein was pooled based on the criteria above, aliquoted and flash-frozen in liquid nitrogen, and stored at −80°C.

#### Inhibitor characterization of 3CLpro enzymes

Inhibition of all 3CLpro enzymes was measured using a continuous, FRET assay. The increase in fluorescence due to the cleavage of a custom-synthesized substrate, UIVT3 (HiLyte Fluor_488_™-ESATLQSGLRKAK-QXL_520_™-NH_2_) (Anaspec) by the enzymes was monitored over time. All K_i_ determinations were performed in Costar 3694 EIA/RIA 96-well, half-area, flat bottom, black polystyrene plates (Corning) using an assay buffer that contained 50mM HEPES (pH 7.5), 0.1mg/mL bovine serum albumin (BSA), 0.01% Triton X-100 and 2mM DTT. The final assay volume was 100μL and all assays were performed in triplicate. 1μL of 100X inhibitor stocks, prepared in 100% DMSO, were added to assay buffer and incubated for 5 minutes at 25°C. Then, 3CLpro enzyme was added to the mixture and incubated for 10 minutes at 25°C. The final concentrations of each coronaviral 3CLpro enzyme are listed below and they are different depending on the coronavirus. All reactions were initiated with the addition of UIVT3 substrate to a final concentration of 2μM.

The fluorescence of the substrate (excitation 485 nm / emission 528 nm) was measured using either a CLARIOstar Plate Reader (BMG Labtech) or Synergy H1 hybrid multi-mode plate reader (Biotek). The initial rates at each inhibitor concentration were obtained by dividing the Relative Fluorescence Units (RFU) produced during the initial rate period of the enzyme by time in minutes, yielding RFU min^−1^. Initial velocities were calculated by dividing the observed velocity in the absence of inhibitor (V_o_) by the initial velocity at different inhibitor concentrations (V_i_). K_i_ values, with the exception of MERS 3CLpro, were obtained by fitting the data to the Morrison equation (Equation 1) for tight-binding inhibitors^45^.

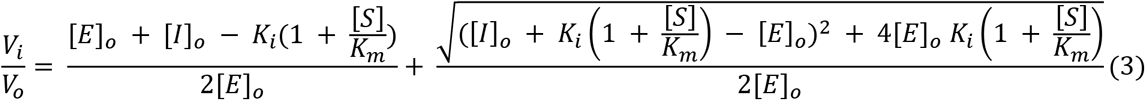

For these fits, the V_max_ was initially set to a value less than 1.2, the UIVT3 substrate concentration was fixed at 2μM, and the K_m_ value was fixed at a value of 250μM. The K_m_ value for the UIVT3 substrate is estimated to be higher as even after correcting for inner-filter effects via dilution experiments, the 3CLpro enzymes were not saturated with substrates as the response of the enzyme to increasing UIVT3 substrate concentrations was linear over the range of 1 to 250μM.

The total enzyme concentrations were experimentally determined by measuring their absorbance at 280nm and using their respective molar extinction coefficients ε_280_ calculated based on their primary amino acid sequences. The values for enzyme concentration [E]_o_ in Equation 3 were not fixed during the curve-fitting process. The resulting enzyme concentrations from the Morrison equation fit are represented in the parentheses following the experimentally determined total enzyme concentrations as follows: 200nM SARS-2 (188 ± 10nM), 200nM SARS (153 ± 2nM), 1000nM MERS, 250nM NL63 (170 ± 4nM), 125nM HKU1 (59 ± 2nM), 100nM HKU4 (60 ± 1nM), 125nM HKU5 (74 ± 1nM), 400nM HKU9 (265 ± 5nM), 200nM 229E (116 ± 3nM), 100nM OC43 (51 ± 1nM), 100nM PEDV (40 ± 1nM), 100nM MHV (75 ± 4nM), 25nM FIPV (36 ± 1nM), 50nM IBV (31 ± 1nM). MERS 3CLpro data were calculated using Equation (4) and fit to the substrate inhibition equation. %V_max_ is the maximum percent activity observed. All fits were performed using GraphPad Prism 8.3.

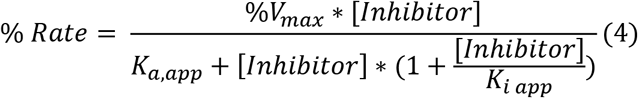

### Mammalian Protease panel

The respective protease in assay buffer (50mM Tris with 100mM sodium chloride and Brij 35 at pH = 8 except for cathepsin D pH = 3.5 and HIV pH = 5.5) was added to assay ready compound plates. The enzymatic reaction was initiated with the addition of indicated substrate in assay buffer. Final concentrations of respective protease and substrate are shown in Table S12. Initial rates were measured by following the fluorescence of the cleaved substrate (Ex/Em 355/460nM) using a Spectramax (Molecular Devices) fluorescence plate reader in the kinetic format.

Percent inhibition values were calculated based on control wells containing no compound (0% inhibition) and wells containing a control compound (100% inhibition). IC_50_ values were generated based on a four-parameter logistic fit model using ActivityBase software (IDBS). Percent activity values were calculated based on control wells containing no compound (100% activity) and wells containing a control compound (0% activity).

### Antiviral activity

The ability of compounds to inhibit viral induced cytopathic effect (CPE) against human coronaviruses (SARS-CoV-1, SARS-CoV-2, hCoV-229E) was assessed by monitoring cell viability using two different assay endpoints in VeroE6 or MRC-5 cells. VeroE6 cells that are enriched for hACE2 expression were batched inoculated with SARS-CoV-2 (USA_WA1/2020) at a multiplicity of infection (MOI) of 0.002 in a BSL-3 lab (Southern Research Institute). Virus inoculated cells are then added to assay ready compound plates at a density of 4,000 cells/well in DMEM containing 2% heat inactivated eFBS. Following a 3-day incubation at 37°C with 5% CO_2_, a time at which virus-induced cytopathic effect is 95% in the untreated, infected control conditions. Cell viability was evaluated using Cell Titer-Glo (Promega), according to the manufacturer’s protocol, which quantitates ATP levels. Cytotoxicity of the compounds was assessed in parallel in assay ready compound plates with non-infected cells.

VeroE6-EGFP cells were seeded at a density of 2,000 cells/well in DMEM containing 2% FCS and 0.08% sodium bicarbonate were seeded into 384-well cell culture microplates containing the serially diluted test compounds. Immediately following cell seeding, virus (SARS-CoV-2 (Belgium strain) or SARS-CoV-1 (CDC 200300592)) was added at a MOI of 0.015 equaling ~30 PFU in a Caps-It isolator system (Rega Institute). The plates were then automatically transferred to an integrated incubator and incubated for 5 days at 37°C with 5% CO_2_, a time at which virus-induced cytopathic effect is 100% in the untreated, infected control conditions (virus controls). Uninfected assay ready compound plates were treated the same way and were used to measure compound cytotoxicity. To monitor the EGFP fluorescence, the plates were transferred to a high-content imager for fluorescence microscopy. A 5x objective was used to capture one field per well (approximately 70% of the well). The images were captured with one channel on auto-focus, a fixed exposure time of 0.023 seconds and with a fixed objective offset (z offset). For the image analysis, the SpotDetector bio-application from the Cellomics software was used, which requires two (2) channels, one (1) for object identification (first channel) and one (1) for signal/spots identification (second channel). The Valid Object Count reported value was the total count of the number of nuclei.

MRC-5 cells, seeded at a density of 20,000 cells/well were incubated overnight in MEM containing 5% FBS at 37oC and 5% CO2. (Wuxi AppTech). The following day test compounds, HCoV-229E virus (ATCC VR-740) (200 TCID50) was added at concentrations which correspond to a multiplicity of infection (MOI) of 0.007, were added to the MRC-5 cells. Cells were incubated for 3 days at 35°C with 5% CO2 and cell viability was evaluated using the CellTiter Glo (Promega) post the 3-day incubation, according to the manufacturer’s protocol. Cytotoxicity of test compounds was assessed in parallel with cells plated 1 day prior to addition of compound, but incubated for 5 days with compound only, and then treated with CellTiter Glo (Promega).

Test compound(s) were tested either alone or in the presence of the P glycoprotein (P-gp) inhibitor, CP-100356 at indicated concentrations of either 0.5 or 2 μM. The inclusion of CP 100356 was to assess if the test compound(s) were being effluxed out of cells due to expression of P-glycoprotein.

Percent effect at each concentration of test compound was calculated based on the values for the no virus control wells and virus containing control wells on each assay plate. The concentration required for a 50% response (EC_50_) value was determined from these data using a 4-parameter logistic model. EC_50_ curves were fit to a Hill slope of 3 when >3 and the top dose achieved ≥ 50% effect. If cytotoxicity was detected at greater than 30% effect, the corresponding concentration data was eliminated from the EC_50_ determination. For cytotoxicity plates, a percent effect at each concentration of test compound was calculated based on the values for the cell only control wells and hyamine or no cell containing control wells on each assay plate. The CC_50_ value was calculated using a 4-parameter logistic model. A therapeutic index (TI) was then calculated by dividing the CC_50_ value by the EC_50_ value.

Drug combination studies were performed using HeLa-ACE2 cells in a high content imaging assay. HeLa-ACE2 cells were seeded at a density of 1.0×103 cells per well in DMEM containing 2% FBS into the 384-well μclear-bottom assay ready compound plates. Plated cells were transferred to the BSL-3 facility where 13μL of SARS-CoV-2 (strain USA-WA1/2020, propagated in Vero E6 cells) diluted in DMEM with 2% FBS was added per well at a concentration to achieve ~30 – 50% infected cells. Plates were incubated for 24 hours at 34℃ with 5% CO2, and then fixed with 25μL of 8% paraformaldehyde for 1 hour at 34℃ with 5% CO2. Plated cells were transferred to the BSL-3 facility where SARS-CoV-2 (strain USA-WA1/2020) was added per well at a concentration to achieve ~30 – 50% infected cells. Plates were incubated for 24 hours at 34℃ with 5% CO2, and then fixed with 25μL of 8% paraformaldehyde for 1 hour at 34℃ with 5% CO2. Plates were washed with 1X phosphate buffered saline (PBS) containing 0.05% Tween 20 after fixation, primary and secondary antibody staining. Human polyclonal sera, from two different recovered patients, was diluted 1:500 in Perm/Wash buffer, added to the plate and incubated at room temperature for 2 hours for primary staining. Six μg/mL of goat anti-human H+L conjugated Alexa 488 together with 8μM of antifade-4,6-diamidino-2-phenylindole dihydrochloride (DAPI) in SuperBlock T20 (PBS) blocking buffer was added and incubated at room temperature for 1 hour in the dark for secondary staining. Plates were imaged using the ImageXpress Micro Confocal High-Content Imaging System (Molecular Devices) with a 10× objective, with 4 fields imaged per well. Images were analyzed using theMulti-Wavelength Cell Scoring Application Module (MetaXpress). The total number of cells were determined with DAPI staining to identify the host-cell nuclei and the number of SARS-CoV-2 infected cells were determined by the SARS-CoV-2 immunofluorescence.

Data were analyzed using Genedata Screener, Version 17.0.1-Standard. Primary invitro screen and the host cell toxicity screen data [% Positive W2 (MultiWaveScoring)] were uploaded to Genedata Screener. For EC_50_ determinations, data were normalized to negative inhibition controls (DMSO) minus positive viral inhibitor controls (2.5μM remdesivir) and for host cell cytotoxicity in the viral infected cells (concentration required for 50% cytotoxicity; CC_50_), data were normalized to negative cytotoxic controls (DMSO) minus positive cytotoxic controls (10μM puromycin). The “% Positive W2 (MultiWaveScoring)” and “Total Cells (MultiWaveScoring)” values were used for analysis of antiviral effect and host cell toxicity, respectively. Compounds were tested in technical triplicates on different assay plates and dose curves were fitted with the four parameter Hill Equation. Replicate data were analyzed using median condensing. GeneData Screener was used to assess drug combination effects, at 90% effect, using the Loewe, Bliss, and HSA models or SynergyFinder using the ZIP model^46^.

In general, a synergy score of >1 and a combination index of <1 indicate that the combination treatment has a synergistic effect^24^. To assess whether synergy could be achieved at high inhibition levels, the isobologram level was set at 0.9 to capture meaningful synergy with a 90% viral reduction (equivalent to a 1log_10_ reduction).

### In vivo efficacy studies in MA-SARS-CoV-1 mouse model

Animal treatment and subsequent observations of animal weight, and virus titration were conducted in the BSL-3 laboratory at University of Maryland College Park under IACUC approved protocols. Mice housed in the BSL-3 were anesthetized with ketamine/xylazine mixture and intranasally inoculated with 1×10^5^ pfu of MA15-SARS-CoV-1 in 50 μL of PBS total volume. In all experiments the vehicle for PF-00835231 was 0.05% (v/v) polysorbate-80, 0.1% (w/v) sodium carboxymethylcellulose, dosed at 10 mL/kg S.C. BID. While on study, mice were weighed daily and observed for clinical signs of disease. At day 4 post-infection, 5 mice per group were euthanized by isoflurane inhalation and lung lobes were either fixed in 4% paraformaldehyde (PFA) at 4°C or placed in 1ml PBS with sterile glass beads at −80°C.

For histopathology, lungs in 4% PFA were removed from the BSL3 at least 24 hours after they were added to PFA and embedded in paraffin for sectioning and H&E staining at the University of Maryland at Baltimore Histology Core. Slides were read by Norimitsu Shirai in the Pfizer Cambridge, DSRD Global Pathology group.

MA15-SARS-CoV-1 lung titers were quantified by homogenizing mouse lungs in 1ml phosphate buffered saline (PBS) using 1.0 mm glass beads and a Beadruptor. VeroE6 cells are plated in 6 well plates with 1×10^5^ cells per well. MA15-SARS-CoV-1 virus titer in plaque forming units was determined by plaque assay: 25 μL of the lung homogenate was added to 225 μL of PBS and diluted 10-fold across a 6-point dilution curve with 200 μL of diluent added to each well. After 1 hour, a 3 mL agar overlay containing MEM was added to each well. Plates were incubated for 3 days at 37°C (5% CO_2_) before plaques were counted.

### LogD, solubility and pKa measurements

The LogD of PF-00835231 was measured at pH 7.4 using the previously described shake-flask method^47^. The LogD of PF-07304814 was predicted using ACDlabs (v2019.1.1). Solubility was measured at room temperature by slurrying the solid drug substance in deionized water and pH buffers for 48 hours, filtering the slurry through 1 0.2μm and analyzing the filtrate using LC-UV spectrometry. The pKa was measured by calculating inflection points in the chemical shift of 1H using solution NMR as a function of pH using ^1^H/^31^P NMR data collected on a 600MHz spectrometer at 298K as described in a previously published method^48^.

### Metabolism of PF-00835231 in human liver microsomes: intrinsic clearance and effects of CYP3A inhibitors

*In vitro* lability in pooled human liver microsomes was determined as follows. PF-00835231 (1 μM) was incubated with human liver microsomes (2.0mg/mL; custom mixed sex pool from 50 donors, Xenotech, Lenexa, KS) in a volume of 1mL potassium phosphate (100mM; pH 7.5) containingmgCl_2_ (3.3mM) and NADPH (1.3mM) at 37°C in a shaking water bath. At 0, 5, 10, 20, 30, 45, and 60 minutes, aliquots (0.1mL) were removed and added to 0.5mL acetonitrile containing saquinavir (0.2 μM) as an internal standard to terminate the reaction. Precipitated protein was removed by spinning in a centrifuge (1700 x g; 5 minutes) and the supernatant was removed in a vacuum centrifuge. The residues were reconstituted in 0.05mL of 1% formic acid in 20% acetonitrile for analysis by high-performance liquid chromatography-tandem mass spectrometry (HPLC-MS). Intrinsic clearance by substrate depletion was calculated as previously described^49^. Enzyme kinetics were determined by product formation. PF-00835231 (0.25-500 μM) was incubated in pooled human liver microsomes (0.5mg/mL) in 0.1mL potassium phosphate (100mM; pH 7.5) containingmgCl_2_ (3.3mM) and NADPH (1.3mM). Incubations were carried out at 37°C by shaking in a humidified incubator (85% relative humidity) for 20 minutes and terminated with the addition of 0.5mL acetonitrile containing saquinavir (0.1 μM) as an internal standard. Terminated incubation mixtures were processed as described above and analyzed by HPLC-MS. Quantitation of the four main metabolites, which had been generated by biosynthesis and qNMR using a previously described method^50^, was done against standard curves ranging from 1-10000nM. Experiments to evaluate the role of CYP3A were conducted in a similar fashion, using a substrate concentration of 4.0 μM, and concentrations of ketoconazole ranging from 1-10000nM. To evaluate the potential contribution of CYP3A5, the comparative effects of ketoconazole vs cyp3cide (both at 1.0 μM) on the formation of four metabolites of PF-00835231 (at 4 μM) were measured in a custom pool of liver microsomes from four CYP3A5 EM donors, using the same incubation conditions described above. All measurements were made in a minimum of triplicate incubations.

### Metabolic stability of PF-07304814 in human liver S9

Human liver S9 (HLS9) was purchased from BioIVT (Westbury, NY) as a custom pool of 6 donors prepared in the absence of EDTA and PMSF. The experimental conditions used for enzyme kinetics studies in liver S9 were chosen to yield linear reaction velocities as determined from preliminary range finding experiments. Stock solutions of PF-07304814 were prepared in 10/90 acetonitrile/water at 10-times the intended incubation concentration. Incubations were conducted in 100mM Tris buffer (pH 7.5) containing 5mMmgCl_2_ and 0.03mg/mL S9 protein at 37°C open to air. The concentrations of PF-07304814 ranged from 0.3 to 300 μM and the final incubation volume was 20 μL. Reactions were initiated by the addition of substrate into enzyme matrix and were terminated after 20 minutes with 100 μL of acetonitrile containing 1% formic acid and IS (50 ng/mL indomethacin). To minimize saturation of MS detection, a 10-fold dilution of samples containing the highest substrate concentrations was conducted prior to precipitation. Samples were vortexed, centrifuged, and 75 μL of supernatant was combined with 200 μL of 1% formic acid in water for injection onto LC-MS system for analysis. Incubations for enzyme kinetic determination in liver S9 were conducted in triplicate. Enzyme kinetics were analyzed as described above for HLM using the Michaelis-Menten model with a second unsaturable component. The apparent intrinsic clearance for PF-07304814 in liver S9 across species was calculated using equation 5.

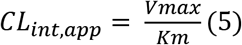

### Reversible inhibition of CYP enzymes in human liver microsomes

CYP selective marker substrates were purchased from various commercial sources: furafylline, ticlopidine, paroxetine, phenacetin, acetaminophen, diclofenac, dextromethorphan and dextrophan were obtained from Sigma Aldrich (St. Louis, MO); S-mephenytoin from Toronto Research Chemicals (North York, ON); verapamil from Sequoia Research Products (Pangbourne, United Kingdom); amodiaquine from Fluka (Bucks, Switzerland); 4’-OH-S-mephenytoin and [2H3]4’-OH-S-mephenytoin from Syncom (Groningen, Netherlands); N-desethylamodiaquine, [2H5]N-desethylamodiaquine, tienilic acid, and [2H3]-dextrophan from Cerilliant (Round Rock, Texas); gemfibrozil glucuronide, 4ˈ-OH diclofenac, and [13C6]4ˈ-OH diclofenac were from Pfizer Inc (Groton, CT). Reversible inhibition was measured in a co-incubation of PF-00835231 with marker substrate (at approximately KM concentration) with HLM (0.01-0.1mg/ml) and NADPH (1.3mM) in 100mM potassium phosphate buffer containing 3.3mMmgCl2 (pH 7.4) in a final incubation volume of 0.2mL in a 37°C dry heat bath open to air. After a 4-20 min incubation time, reactions were quenched with 2-volumes of acetonitrile containing internal standard (2H or 13C labeled analyte). To qualitatively determine the potential for TDI, all incubation components except marker substrate were preincubated for 30 min, followed by addition of marker substrate for the specified reaction time. PF-00835231 was tested at concentrations 0.1 to 200 μM in triplicate. Terminated incubation mixtures were centrifuged at 2000 g for 5 min, resulting supernatants were evaporated to dryness under nitrogen followed by reconstitution in mobile phase (20% acetonitrile in 1% aqueous formic acid). Samples were analyzed by LC-MS. Analytes were quantified versus a standard curve using GraphPad Prism v8 (San Diego, CA) or Sciex Analyst software. IC_50_ calculations were conducted in GraphPad using equation 6.

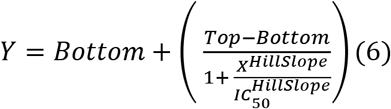

TDI signal was defined having either at least a 1.5-fold decrease in IC_50_ value after a 30-minute preincubation or there was at least a 20% increase in inhibition observed after a 30-minute preincubation for any concentration examined. A TDI signal resulted in determination of K_I_ and k_inact_. For inhibition studies conducted in cassette format, incubations were conducted as described above with the following modifications: HLM concentration was 0.03mg/mL, probe substrates bupropion and testosterone were excluded, and the activity reaction time was 6 minutes.

### Time-dependent inhibition of CYP enzymes in human liver microsomes

K_I_ and k_inact_ studies were measured in a co-incubation of PF-00835231 with HLM (0.2-0.3mg/mL) and NADPH (1.3mM) in 100mM potassium phosphate buffer containing 3.3mMmgCl_2_ (pH 7.4) in a final incubation volume of 0.2mL in a 37°C dry heat bath open to air. After a 1-50-minute preincubation time, an aliquot of incubate was diluted 20-fold into prewarmed buffer containing marker substrate (5-10 times above K_M_ concentration), followed by an activity incubation: 20 μM midazolam (midazolam 1′-hydroxylation) 4 minutes with 0.2mg/mL HLM; CYP3A, 386 μM testosterone (testosterone 6β-hydroxylation) 11 minutes with 0.3mg/mL HLM. Reactions were quenched with 2-volumes of acetonitrile containing internal standard (^2^H or ^13^C labeled marker product) and treated as described above for IC_50_ samples. Estimation of the TDI inactivation parameters was performed using Excel and GraphPad software. Percent activity remaining was obtained by normalizing the concentration of marker substrate formed in each sample to the mean solvent control at the first preincubation time point. The natural log (ln) of the percentage remaining activity was plotted against the preincubation time. The slope (−k_obs_, observed rate) of each line was then calculated for the linear portion of the curve and the non-linear outliers, as determined by GraphPad automatic outlier elimination, were excluded. A statistical test was applied at each inhibitor concentration to evaluate if k_obs_ was significantly different from the solvent control (equation 7)^51^.

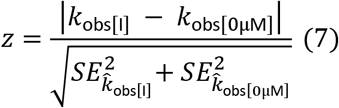

In this equation, k_obs[I]_, k_obs[0 μM]_, and S.E. represent the inactivation rate at each inhibitor concentration, inactivation rate with solvent control, and standard error, respectively. When p<0.05, there was statistically significant or measurable TDI. K_I_ and *k_inact_* were calculated from the nonlinear regression of a 3-parameter Michaelis‑Menten equation (equation 8) using GraphPad.

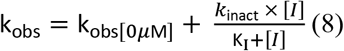

### *In vitro* transporter inhibition studies using transporter-transfected cells

HEK293 cells, wild-type and stably transfected with OATP1B1, OATP1B3, OCT1, OCT2, MATE1 and MATE2K, were seeded at a density of 0.5-0.7 × 10^5^ cells/well on BioCoat™ Poly-D-Lysine 96-well plates (Corning) and grown in Dulbecco’s modified Eagle’s medium containing 10% FBS, 1% sodium pyruvate, 1% GlutaMAX™, 1% Gentamicin and 1% nonessential amino acids for 48-72 hours at 37°C, 90% relative humidity, and 5% CO_2_. For the inhibition studies, HEK293 cells were washed three times with warm transport buffer (Hanks’ balanced salt solution with 20mM 4-(2-Hydroxyethyl) piperazine-1-ethanesulfonic acid, pH 7.4) followed by incubation with test compounds containing probe substrates: 10 or 20 μM [^14^C]-metformin (OCT1, OCT2, MATE1, MATE2K), 0.5 μM [^3^H] para-aminohippuric acid (OAT1), 0.1 μM [^3^H] estrone-3-sulfate (OAT3) or 0.5 μM rosuvastatin (OATP1B1/1B3). Uptake was terminated by washing cells at least three times with ice-cold transport buffer and then lysing with 0.2mL of scintillation fluid or 0.225mL of methanol containing internal standard. For radiolabeled compounds, radioactivity in each sample was quantified by measurement on MicroBeta (Perkin Elmer). For rosuvastatin, cell extracts were dried down under nitrogen and reconstituted in 50:50 (v:v) methanol:water prior to injection onto a LC-MS/MS system. The total cellular protein content was determined by using a BCA Protein Assay Kit (Pierce Biotechnology) following the manufacturer’s protocol. Uptake ratio was then derived as a ratio of accumulation in transfected cells to accumulation in wild-type cells. The half-maximal inhibitory concentration (IC_50_) for each transporter was calculated in GraphPad Prism.

(HEK = Human embryonic kidney; OATP = Organic anion transporting polypeptide; OCT = Organic cation transporter; MATE= Multi drug and toxin extrusion protein)

### *In vitro* transporter inhibition studies using membrane vesicles

Vesicle assay buffer (10mM Tris base, 250mM sucrose, 10mM magnesium chloride) and stop buffer (assay buffer plus 100mM sodium chloride) were prepared at pH 7.4. For inhibition assays, HEK293, human BCRP and human MDR1 (P-gp) vesicles were diluted to 50 μg/well in assay buffer and were treated with 5mM ATP, 0.2 μM rosuvastatin (BCRP), or 0.2 μM N-methyl quinidine (MDR1) and varied concentrations of test compounds. BCRP and MDR1 assays were incubated for 1 and 1.5 minutes, respectively, at 22°C while shaking. Reactions were stopped by adding 0.2mL of ice-cold stop buffer. The entire reaction was quickly removed from the assay plate and filtered on Multiscreen filter plate (Millipore) and then washed four times with ice-cold stop buffer. Intravesicle samples were extracted by treating the vesicles with 0.15mL of methanol containing internal standard. Vesicles were shaken for 25 minutes at room temperature. Vesicle extracts were transferred to 96-well polypropylene deep-well plates by centrifugation and dried down under nitrogen. Samples were reconstituted in 50:50 (v:v) methanol:water prior to injection onto a LC-MS/MS system. Percent activity values were exported into GraphPad Prism to estimate IC_50_ values.

(MDR1 = Multi drug resistant AKA P-glycoprotein; BCRP = Breast cancer resistant protein)

### Plasma protein, liver microsome, and liver S9 binding

Frozen plasma in K_3_EDTA was purchased from BioIVT and Dulbecco’s phosphate buffered saline (DPBS) and HCl were purchased from Sigma. Fraction unbound was determined by equilibrium dialysis using an HTD 96 device (HTDialysis, LLC, Gales Ferry, CT) assembled with 12-14k MWCO membranes. Plasma was thawed and adjusted to pH 7.4 with 1N HCl prior to use. Dialysis chambers were loaded with 150 μL plasma and 150 μL PBS in the donor and receiver chambers, respectively. Human liver microsomes were diluted to a concentration of 0.8mg/mL in 100mM potassium phosphate buffer, donor chambers were loaded with 150 μL diluted HLM and receiver chambers 150 μL potassium phosphate buffer. Human liver S9 was diluted to a concentration of 0.03mg/mL in 100mM potassium phosphate buffer, donor chambers were loaded with 150 μL diluted HLS9, and receiver chambers 150 μL potassium phosphate buffer. The dialysis plate was sealed with a gas-permeable membrane and stored in a 37°C water-jacked incubator maintained at 75% relative humidity and 5% CO_2_, on a 100 rpm plate shaker. After a 6-hour incubation, samples were matrix-matched and quench by protein precipitation, followed by LC-MS analysis. A set of satellite samples was included to measure stability after a 6-hour incubation. Incubations were conducted with 4 to 12 replicates. Fu was calculated by dividing the analyte-to-internal standard peak area ratio or analyte concentration in the buffer sample by the signal in the donor sample, corrected for any dilution factors. The HLM binding measured at 1mg/mL was adjusted to account for the HLM protein concentration (2mg/mL) using a dilution equation^52^. All incubations had >70% analyte recovery and >70% stability in 6 hours.

### Preclinical pharmacokinetic studies

All activities involving animals were carried out in accordance with federal, state, local and institutional guidelines governing the use of laboratory animals in research in an AAALAC accredited facility and were reviewed and approved by Pfizer’s Institutional Animal Care and Use Committee.

Rat PK studies were done at Pfizer (Groton, CT) or BioDuro Pharmaceutical Product Development Inc. (Shanghai, PRC); Jugular vein-cannulated male Wistar-Hannover rats were purchased from Charles River Laboratories, Inc. (Wilmington, MA) or Vital River (Beijing, China) and were typically 7-10 weeks of age at the time of dosing. During the pharmacokinetic studies all animals were housed individually. Access to food and water was provided ad libitum. Compounds were administered i.v. via the tail vein (n = 2 or 3), dosed as a 1mg/ml solution using standard compatible excipients (PF 00835231, 2mL/kg or PF 7304814, 1mL/kg) for a resulting dose of 2mg/kg PF 00835231 or 1.17mg/kg PF 07304814. Serial blood samples were collected via the jugular vein cannula at predetermined timepoints after dosing. Animals were monitored for pain or distress throughout the study, with at least daily monitoring during normal husbandry prior to study start. At the completion of the study, animals were euthanized by overdose of inhaled anesthesia followed by exsanguination. Blood samples were collected into tubes containing K3EDTA and stored on ice until centrifugation to obtain plasma, which was stored frozen at −20 °C or lower. Urine samples were collected at room temperature and stored frozen at 20°C or lower at the end of each time interval.

Dog PK studies were done at Pfizer (Groton, CT); animal care and in vivo procedures were conducted according to guidelines from the Pfizer Institutional Animal Care and Use Committee. Male Beagle dogs were purchased from Marshall BioResources (North Rose, New York) and were typically 1-5 years of age at the time of dosing. Compounds were administered i.v. via the cephalic vein (n = 2), dosed as a 2mg/ml solution using standard compatible excipients (PF 00835231, 0.5mL/kg or PF 7304814, 0.5mL/kg) for a resulting dose of 1mg/kg PF 00835231 equivalents or 1.17mg/kg PF 07304814. Serial blood samples were collected via the jugular vein at predetermined timepoints after dosing. Animals were monitored for pain or distress throughout the study, with at least daily monitoring during normal husbandry prior to study start. Blood samples were collected into tubes containing K3EDTA and stored on ice until centrifugation to obtain plasma, which was stored frozen at −20°C or lower. Urine samples were collected at room temperature and stored frozen at −20°C or lower at the end of each time interval.

Non-human primate PK studies were conducted at Pfizer (Groton, CT); All procedures performed on the animals were in accordance with regulations and established guidelines and were reviewed and approved by an Institutional Animal Care and Use Committee through an ethical review process. Male Cynomolgus monkeys were purchased from Covance (Princeton, NJ), Charles River Laboratories, Inc. (Wilmington, MA), or Envigo Global Services (Indianapolis, IN); animals 3-8 years of age were used in PK studies. Compounds were administered i.v. via the saphenous or cephalic vein (n = 2), dosed as a 1mg/ml solution using standard compatible excipients (PF 00835231, 1mL/kg or PF 7304814, 0.5mL/kg) for a resulting dose of 1mg/kg PF 00835231 or 1.17mg/kg PF 07304814. PF-00835231 was also administered via oral gavage (n=2) as a 1mg/ml suspension in 0.5% methylcelluose (w/v) in water, at a dose volume of 5mL/kg, for a final dose of 5mg/kg. Serial blood samples were collected via the femoral vein at predetermined timepoints after dosing. Animals were monitored for pain or distress throughout the study, with at least daily monitoring during normal husbandry prior to study start. Blood samples were collected into tubes containing K3EDTA and stored on ice until centrifugation to obtain plasma, which was stored frozen at −20°C or lower. Urine samples were collected at room temperature and stored frozen at −20°C or lower at the end of each time interval.

Plasma and urine samples were processed using protein precipitation with acetonitrile:methanol containing internal standard propranolol (50 ng/ml) followed by quantitation against a standard curve (0.1-2500 ng/ml) prepared in blank plasma. Pharmacokinetic parameters were calculated using noncompartmental analysis (Watson v.7.5, Thermo Scientific). The area under the plasma concentration-time curve from t = 0 to infinity (AUCinf) was estimated using the linear trapezoidal rule. Plasma clearance (CLp) was calculated as the i.v. dose divided by AUCinf. The terminal rate constant (kel) was calculated by linear regression of the terminal phase of the log-linear concentration-time curve and the terminal elimination t1/2 was calculated as 0.693 divided by kel. Apparent steady state distribution volume (Vdss) was determined by clearance multiplied by mean residence time. Percent excreted in urine was calculated as the amount of analyte in the urine divided by the amount dosed. Oral bioavailability was defined as the dose-normalized AUC after oral administration divided by the dose-normalized AUC after i.v. administration. Percent conversion of prodrug to active metabolite was calculated by dividing the dose-normalized PF-00835231 AUCinf after i.v. dosing of prodrug PF-07304814 by the AUCinf after i.v. dosing of PF-00835231 incorporating their respective differences in molecular weight.

### Bioanalytical LC-MS analysis

Metabolic profiling samples were analyzed by ultrahigh-performance liquid chromatography coupled to UV spectrometry and high-resolution mass spectrometry (UHPLC-UV-HRMS). The system consisted of a Thermo Accela quaternary HPLC pump (Waltham, MA), Thermo Accela diode array UV/VIS detector, and CTC Analytics autoinjector (Zwingen, Switzerland), hyphenated with a Thermo Orbitrap Elite high-resolution mass spectrometer. Samples (0.01mL) were injected onto a Phenomenex XB-C18 column (2.1 × 100mM, 2.6 μm) (Torrance, CA) maintained at 45°C, at a flow rate of 0.4mL/min. The mobile phase components were A: 0.1% formic acid in water and B: acetonitrile. The initial mobile phase composition of 5% B was held for 0.5 minutes followed by sequential linear gradients to 70% B at 11 minutes and 95% B at 13 minutes. This composition was held for 1 minute followed by re-equilibration at initial conditions for 1 minute. The eluent was passed through the UV detector (scanning from 200-400nm) before introduction into the mass spectrometer. The mass spectrometer was operated in positive mode with ion source temperatures, potentials, and gas flow settings adjusted to optimize the signal for PF-00835231.

LC-MS analysis of remaining PF-00835231 and PF-07304814 samples was typically performed using tandem liquid chromatography-mass spectrometry (LC-MS/MS) with a Sciex Triple Quad 5500 or 6500 mass spectrometer (Sciex, Framingham, MA), equipped with electrospray sources and Agilent 1290 binary pump (Santa Clara, CA). Aqueous mobile phase (A) was comprised of 1.0% formic acid in water and organic mobile phase (B) consisted of 1.0% formic acid in acetonitrile. Ten μl of sample was injected onto an Acquity UPLC BEH C18 (2.1 × 50mm, 1.7 μm) (Waters Corporation, Milford, MA) or Halo C18 (2.1 × 30mm, 2.7 μm) (Advanced Materials, Wilmington, DE) column at room temperature with a flow rate of 0.5 mL/min. The gradient program began with 5% initial mobile phase B held for 0.3 minutes, followed by a linear gradient to 95% B over 2 minutes, held at 95% B for 0.3 minutes followed by re-equilibration to initial conditions for 0.6 minutes. MS was operated in multiple reaction monitoring (MRM) mode, in positive detection mode, with the following mass transitions and collision energies: PF-00835231 473.2/187.0 (CE 16), PF-07304814 553.1/267 (CE 24), propranolol 260.2/116.2 (CE 30), and indomethacin 358.1/139.0 (CE 27).

CYP inhibition samples were analyzed using MRM in positive ion mode. 1A2 reactions were analyzed for acetaminophen 152.1/110.0 and [^2^H_7_]-acetaminophen 159.0/115.0 (CE 22) using a Phenomonex Synergi Hydro-RP column (2×30mm, 2.5μm). 2B6 reactions were analyzed for hydroxybupropion 256.0/139.0 and [^2^H_6_]-hydroxybupropion 262.0/139.0 (CE 31) using a Halo C18 column (2.1×30mm, 2.7μm). 2C8 reactions were analyzed for N-desethylamodiaquine 328.2/283.0 and [^2^H_5_]-N-desethylamodiaquine 333.2/283.0 (CE 23) using a Phenomenex Synergi Hydro-RP column (2×30mm, 2.5 μm). 2C9 reactions were analyzed for 4′-hydroxydiclofenac 213.0/266.0 and [^13^C_6_]-4′-hydroxydiclofenac 318.0/272.0 (CE 20) using a Halo C18 column (2.1×30mm, 2.7 μm). 2C19 reactions were analyzed for 4′-hydroxymephenytoin 235.2/150.1 and [^2^H_3_]-4′-hydroxymephenytoin 238.1/150.1 (CE 26) using a Halo C18 column (2.1×30mm, 2.7 μm). 2D6 reactions were analyzed for dextrorphan 258.1/201.1 and [^2^H_3_]-dextrorphan 261.0/201.1 (CE 31) using a Halo C18 column (2.1×30mm, 2.7 μm). 3A4 reactions were analyzed for 1′-hydroxymidazolam 342.1/324.0 and [^2^H_4_]-1′-hydroxymidazolam 346.1/328.0 (CE 30) using a Halo C18 column (2.1×30mm, 2.7μm) or 6β-hydroxytestosterone 305.2/269.2 and [^2^H_3_]-6β-hydroxytestosterone 308.2/272.7 (CE 25) using a Halo C18 column (2.1×50mm, 2.0μm).

Analyst software was used to measure peak areas and peak area ratios of analyte to internal standard were calculated. A calibration curve was constructed from the peak area ratios (analyte to internal standard) with a weighted linear (1/x^2^) regression, from which unknown samples concentrations were calculated.

### Human pharmacokinetic predictions

#### Scaling in vitro human liver microsomal or liver S9 apparent CLint to in vivo CLint

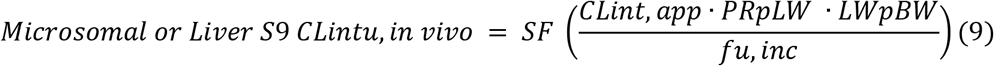

CLint, app = apparent intrinsic clearance, PRpLW = Microsomal or S9 protein per liver weight (45 or 121mg/g respectively), PRpLW = liver weight per bodyweight (21g/kg), fu,inc = HLM or S9 unbound fraction, SF = Pfizer scaling factor (x1.3 for HLM, x1 for S9)

#### Scaling in vivo CLint to human plasma clearance

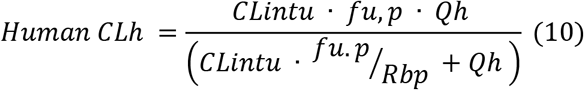

CLintu = scaled *in vivo* intrinsic clearance, fu,p =plasma fu, Qh = hepatic blood flow (20 mL/min/kg), Rbp = *in vitro* blood to plasma ratio

#### Prediction of PF-00835231 human Vdss

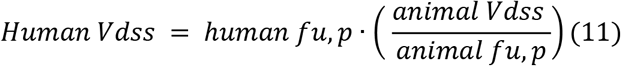

#### Calculation of predicted human half-life

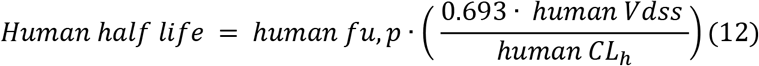

#### Calculation of human dose

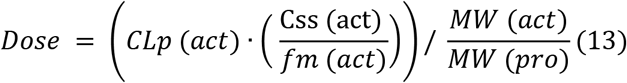

Rearranged from published equation factoring in MW differences^53^. act = PF-00835231, pro = PF-07304814, CLp = plasma clearance, Css = steady state concentration, fm = %conversion from PF-07304814 to PF-00835231, MW = molecular weight (PF-07304814 = 552.5, PF-00835231 = 472.5)

### Physiologically-based pharmacokinetic (PBPK) modeling of PF-00835231

A commercially available dynamic PBPK model, Simcyp population-based simulator (version 18.2; Certara UK Limited, Simcyp Division, Sheffield, United Kingdom), was used in the present study^54^. Physicochemical and pharmacokinetic parameters of PF-00835231 for the PBPK models are summarized in Table S10. Since PF-07304814 (prodrug) was predicted to be converted to PF-00835231 rapidly and extensively *in vivo*, the simulation was performed assuming an intravenous infusion of PF-00835231 with the conversion efficiency of 75% from PF-07304814, which was predicted from animal data. For the prediction of DDIs of PF-00835231 with itraconazole, the vendor-verified compound files in Simcyp library were used, i.e., itraconazole (sv-itraconazole_fed capsule) with competitive K_i_ = 0.0013 μM and itraconazole metabolite (sv-OH-itraconazole) with competitive K_i_ = 0.0023 μM.

Simulation of clinical trials was performed with a virtual population of healthy volunteers in 10 trials of 10 subjects (total 100 subjects), each aged 20 to 50 years with a female/male ratio of 0.5, whose CYP3A4 degradation rate constant (*k_deg_*) was 0.019 h^−1^ in liver and 0.030 h^−1^ in intestine. The output sampling interval in Simcyp simulation toolbox was set to 0.2 hours in all simulations. To predict DDIs of PF-00835231 with itraconazole, PF-00835231 at 320mg/day (equivalent to PF-00835231 formed following PF-07304814 ~500mg/day taking into account conversion and molecular weight differences) was administered IV for 10 days (days 5 to 15) to a virtual population with and without 15-day repeated oral administration of itraconazole 200mg once daily (days 1 to 15). Pharmacokinetic parameters such as maximal plasma concentration (C_max_), area under the plasma concentration time-curve from time zero to 24 hours post dose (AUC) and the ratios of C_max_ (C_max_R) and AUC (AUCR) in treatment groups relative to control groups were obtained from Simcyp outputs.

The model-predicted C_max_R and AUCR for PF-00835231 were ~2x at the daily dose of 320mg/day (corresponding to ~500mg/day PF-07304814) (Table S11).

### Ceff projection of protease inhibitor to the clinic

The inhibitory quotient (IQ) has been a useful metric for translating preclinical antiviral potencies to the clinic across a number of viral diseases as indicated in the FDA guidance^55^. IQ is defined as the human C_min,u_ unbound concentration divided by the *in vitro* unbound (serum adjusted) EC_50,u_ value in the antiviral assay (equation 14).

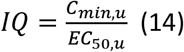

Some antiviral therapies have shown significant benefit with IQ close to 1^56^; however, rapidly controlling viral replication frequently requires maintaining an exposure at least 10x higher than *in vitro* EC_50_^30^. Clinically approved protease inhibitors have effectively decreased viral loads when dosed at IQ values from 1-100, when protein binding and site of action exposure are taken into account^30^. Importantly, antivirals in general and, specifically, protease inhibitors can potentially lead to increased mutations and additional drug resistance when dosed at an IQ less than 1^57^.

How high an IQ value is required depends on the steepness of the dose response curve. The hill coefficient (m), and the EC_50_ are related to the *in vitro* antiviral activity at a range of concentrations (C) by equation 15:

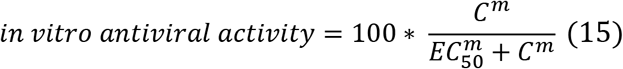

PF-00835231 shows a high hill coefficient (m=3) across a range of *in vitro* antiviral assays, like those of clinical protease inhibitors targeting HIV and HCV^21,58^. There is only a 2- to 3-fold difference between the antiviral EC_50_ and EC_90_ concentrations (Fig. 2), rather than the typical 9-fold difference for antiviral agents with hill coefficients of 1. Therefore, relatively small ratios of exposure to EC_50_ values (3-10) are related to near complete viral suppression.

### Rat 24-hour continuous intravenous infusion GLP toxicity study

The study was conducted by Charles River Laboratories for Pfizer Inc, in accordance with the U.S. Department of Health and Human Services, Food and Drug Administration, United States Code of Federal Regulations, Title 21, Part 58: Good Laboratory Practice for Nonclinical Laboratory Studies. Briefly, 8-week old male and female Sprague-Dawley rats (n=15 per sex per group) implanted with femoral catheters exteriorized between the scapulae were assigned to vehicle, or PF-07304814 dose groups (80, 360 and 1000mg/Kg). The vehicle and test article were administered once on Day 1 as a 24-hour continuous IV infusion at a rate of 2.5 mL/Kg/hour. Main group rats (n=10 per sex per group) were euthanized on Day 2 and recovery/delayed toxicity group (n=5 per sex per group) on Day 14. Study evaluations included detailed clinical observations, body weights, food consumption, functional observation battery, clinical pathology, toxicokinetics, and gross and microscopic pathology.

### Rat 4-day continuous intravenous infusion, exploratory nonGLP toxicity study

The study was conducted with Pfizer Worldwide R & D, and due to the exploratory nature of this study, compliance with GLP regulations was not required. Male Sprague-Dawley rats (n=3 per group) were administered either vehicle or PF-00835231 (at 24.6 and 246mg/kg/day) by continuous intravenous infusion via a femoral venous catheter for 4 days, followed by necropsies on Day 5. Study evaluations included clinical observations, body weights, clinical pathology, toxicokinetics, gross pathology and microscopic pathology (control and high dose groups only).

### Bacterial reverse mutation assays

PF-00835231 and PF-07304814 were tested to evaluate their mutagenic potential by measuring their ability to induce reverse mutations at selected loci of several strains of *Salmonella typhimurium* and at the tryptophan locus of *E. coli* strain WP2 *uvr*A in the presence and absence of an exogenous metabolic activation system using standard protocols^59–61^. These GLP studies were conducted by BioReliance Corporation for Pfizer Inc.

### Effect of PF-00835231 and PF-07304814 on hERG potassium channels

The *in vitro* effects of PF-00835231 and PF-07304814 on the hERG (human ether-à-go-go-related gene) channel current (a surrogate for *I_Kr_*, the rapidly activating delayed rectifier cardiac potassium current ^62^ were tested using standard protocols. This channel has been selected for evaluation because inhibition of *I_Kr_* is the most common cause of cardiac action potential prolongation by non-cardiac drugs ^63–65^. Increased action potential duration causes prolongation of the QT interval and has been associated with a dangerous ventricular arrhythmia, *torsade de pointes*^63^. The concentration-response relationship of the effect of PF-00835231 and PF-07304814 on the hERG potassium channel current was evaluated at near-physiological temperature in stably transfected mammalian cells that express the hERG gene using standard protocols. These GLP studies were conducted by Charles River Laboratories for Pfizer Inc.

### Secondary/off-target pharmacology

Secondary pharmacology studies were conducted by Eurofins Cerep on behalf of Pfizer Inc. The in vitro off-target pharmacology of PF-07304814 and PF-00835231 was individually assessed at 100 μM in a broad target profiling panel which represents targets with known links to potential safety concerns and includes G-protein coupled receptors, ion channels, transporters, and enzymes according to established protocols.

### Hemocompatibility studies

The effects of PF-07304814 and PF-00835231 on human red blood cell hemolysis and plasma flocculation were evaluated in GLP-compliant studies conducted by Charles River Laboratories on behalf of Pfizer Inc. Briefly, whole blood samples from human volunteers were incubated with a range of concentrations of PF-07304814 and PF-00835231 (in relevant vehicles) and hemolysis was evaluated by determination of whole blood hematocrit, whole blood hemoglobin concentration, plasma hemoglobin concentration, plasma hemolytic index, and visual macroscopic hemolysis assessment. Flocculation was evaluated by determination of the plasma turbidity index and visual flocculation assessment.

---

**Fig. S1.**
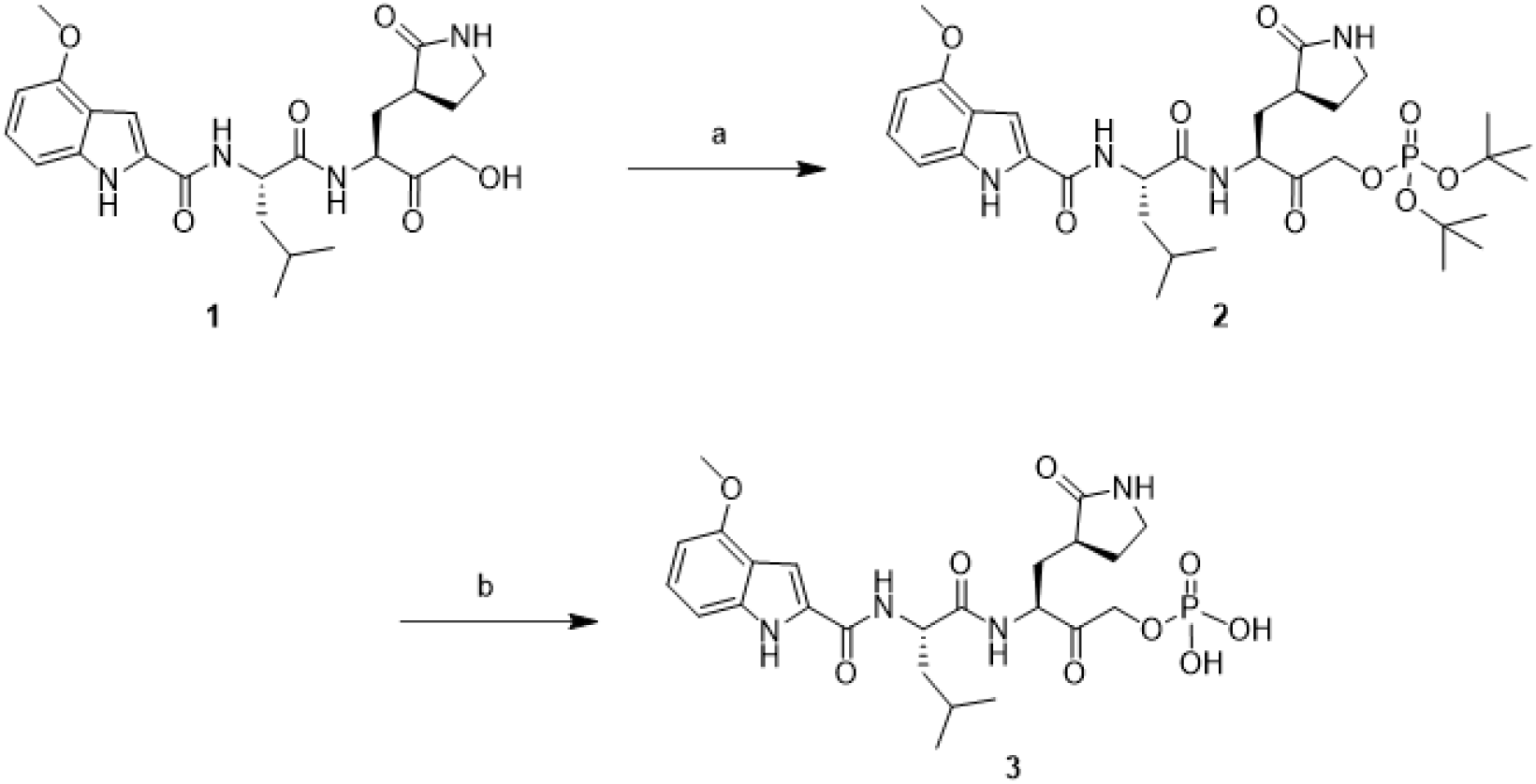
Synthesis of PF-07304814 and PF-00835231 with reagents and conditions. **a**) Di-tert-butyl N,N-dipropan-2-ylphosphoramidite, tetrazole, tetrahydrofuran, 0°C to room temp., then H2O2, 0°C, 91% over 2 steps; **b**) CF3COOH, CH2Cl2, 0°C, 54%.

**Fig. S2.**
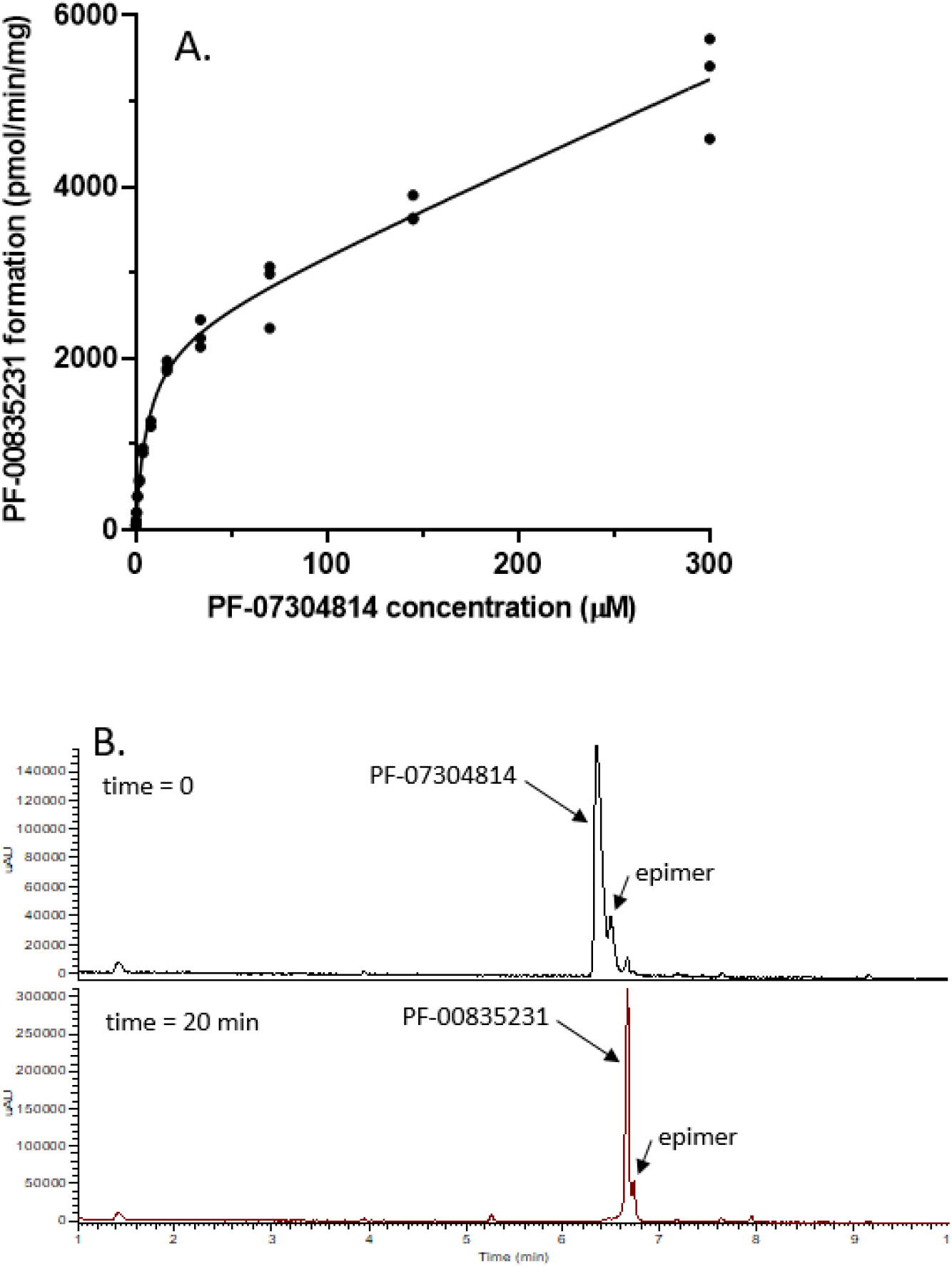
Metabolism of PF-07304814 in human liver S9. (**A)** Substrate saturation plot of the metabolism of phosphate prodrug PF-07304814 to the active drug PF-00835231 in human liver S9 fraction. (**B**) HPLC-UV chromatograms of extracts of an incubation of phosphate prodrug PF-07304814 (Rt 6.3 min) in human liver S9 demonstrating complete conversion to the active entity PF-00835231 (Rt 6.7 min).

**Table S1.** Summary of the In Vitro Antiviral Activity for PF-00835231 on Related Coronaviruses in Vero E6 cells with Efflux Inhibitor.

| **Drug Treatment** | **EC<sub>50</sub>**<br>**SARS-CoV-2**<br>**μM ± Std. Dev** | **EC<sub>50</sub>**<br>**MERS**<br>**μM ± Std. Dev** | **EC<sub>50</sub>**<br>**SARS-CoV-1**<br>**μM ± Std. Dev** | **EC<sub>50</sub>**<br>**MA-15**<br>**μM ± Std. Dev** |
| --- | --- | --- | --- | --- |
| PF-00835231<br>+ 2 μM EI | 0.06 ± 0.03 | 0.04 ± 0.01 | 0.09 ± 0.04 | 0.08 ± 0.01 |

EI = efflux inhibitor (CP-00100356); Std. Dev. = standard deviation; N=3 for SARS-CoV-2 and MERS: N=2 for SARS-CoV-1 and MA-15

**Table S2.**
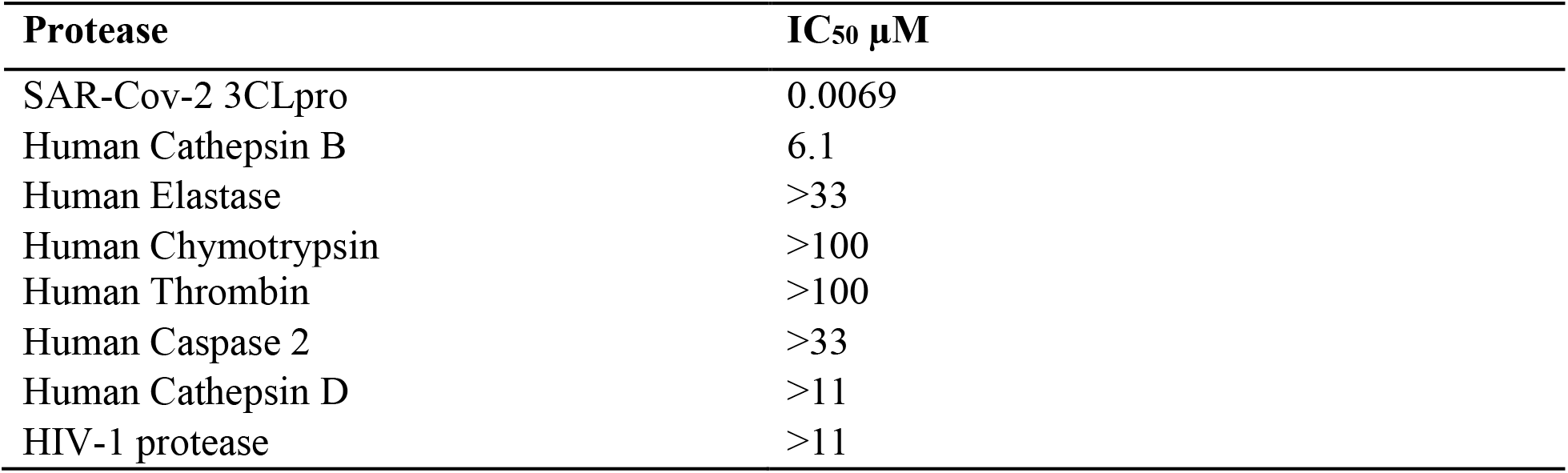
Activity of PF-00835231 against human proteases and HIV protease.

**Table S3.** Evaluation of the fractional metabolism of PF-00835231 in human liver microsomes and recombinant CYP3A using selective CYP3A inhibitor ketoconazole.

| **Metabolite** | **K<sub>M</sub> (μM)** | **V<sub>max</sub>**<br>**(pmol/min/mg)** | **CL<sub>int</sub>**<br>**(μL/min/mg)** | **f<sub>CL</sub>** | **% Inhibition**<br>**by**<br>**ketoconazole** | **f<sub>m</sub>**<br>**(CYP3A)** |
| --- | --- | --- | --- | --- | --- | --- |
| Metabolite 1 | 124 | 321 | 2.6 | 0.61 | 86 | 0.53 |
| Metabolite 2 | 119 | 91 | 0.76 | 0.18 | 90 | 0.16 |
| Metabolite 3 | 120 | 68 | 0.57 | 0.13 | 82 | 0.11 |
| Metabolite 4 | 43 | 12 | 0.30 | 0.07 | 82 | 0.06 |
| **Sum** |  |  | **4.2** |  |  | **0.86** |

CL<sub>int</sub> = Intrinsic clearance; CYP = Cytochrome P450; f<sub>CL</sub> = Fractional clearance; f<sub>m</sub> = Fraction metabolized; HLM = Human liver microsomes; K<sub>M</sub> = Concentration at 50% maximum velocity; rCYP = Recombinant human CYP; V<sub>max</sub> = Maximum initial velocity.

**Table S4.** Reversible CYP inhibition by PF-07304814 or PF-00835231 in human liver microsomes using individual CYP substrates in the presence of NADPH.

| CYP | Enzyme Reaction | Zero-Minute<br>Preincubation (T<sub>0</sub>)<sup>a</sup> |  | 30-Minute Preincubation<br>(T<sub>30</sub>)<sup>a</sup> |  |
| --- | --- | --- | --- | --- | --- |
|  |  | PF-07304814 | PF-00835231 | PF-07304814 | PF-00835231 |
|  |  | IC<sub>50</sub> (μM) | IC<sub>50</sub> (μM) | IC<sub>50</sub> (μM) | IC<sub>50</sub> (μM) |
| 1A2 | Phenacetin<br>O-dealkylation | >100 | >200 | >100 | >200 |
| 2B6 | Bupropion<br>hydroxylation | ND | >200 | ND | >200 |
| 2C8 | Amodiaquine N-dealkylation | >100 | >200 | >100 | >200 |
| 2C9 | Diclofenac<br>4'-hydroxylation | >100 | >200 | >100 | >200 |
| 2C19 | S-Mephenytoin<br>4'-hydroxylation | >100 | >200 | >100 | >200 |
| 2D6 | Dextromethorphan<br>O-demethylation | >100 | >200 | >100 | >200 |
| 3A4/5 | Midazolam<br>1'-hydroxylation | >100 | >200 | >100 | >200 |
| 3A4/5 | Testosterone 6β-hydroxylation | ND | 108 (70-172) | ND | 62.4 (41-98) |

Confidence interval shown in parenthesis; CYP = Cytochrome P450; IC<sub>50</sub> = 50% inhibitory concentration; NADPH = Reduced form of nicotinamide adenine dinucleotide phosphate; NC = Not calculated; ND = Not determined, TDI = Time-dependent inhibition; T<sub>0</sub> = Time zero; T<sub>30</sub> = Time 30 minutes.
<sup>a</sup>Average data obtained from triplicate samples for each test article concentrations were used to calculate IC<sub>50</sub> values.

**Table S5.** Time dependent CYP3A4/5 inhibition by PF-00835231 in human liver microsomes using individual CYP substrates.

| **CYP** | **Probe Substrate**<br>**(Concentration)** | **$k_{\text{inact}} \pm \text{SE}$**<br>**($\text{min}^{-1}$)** | **$K_{\text{I}} \pm \text{SE}$**<br>**($\mu\text{M}$)** | **$k_{\text{inact}} / K_{\text{I}}$**<br>**($\text{mL}/\mu\text{mol}/\text{min}$)** |
| --- | --- | --- | --- | --- |
| 3A4/5 | Midazolam (20 $\mu\text{M}$) | $0.0300 \pm 0.0020$ | $163 \pm 31$ | 0.184 |
| 3A4/5 | Testosterone (386 $\mu\text{M}$) | $0.0434 \pm 0.0007$ | $168 \pm 8$ | 0.258 |

CYP = Cytochrome P450; HLM = Human liver microsomes; $K_{\text{I}}$ = Apparent inactivation constant at half-maximal rate of inactivation; $k_{\text{inact}}$ = Maximal rate of enzyme inactivation; $k_{\text{inact}}/K_{\text{I}}$ = Measure of inactivator efficiency; NADPH = Reduced form of nicotinamide adenine dinucleotide phosphate; SE = Standard error; TDI = Time-dependent inhibition.

**Table S6.** *In vitro* transporter inhibition by PF-07304814 or PF-00835231 using probe substrates.

| Transporter | Probe Substrate (Concentration) | IC<sub>50</sub> (μM) |  |
| --- | --- | --- | --- |
|  |  | PF-07304814 | PF-00835231 |
| MDR1/P-gp | N-methyl quinidine (0.2 μM) | >300 | 65.6 |
| BCRP | Rosuvastatin (0.2 μM) | 238 | 19.5 |
| OATP1B1 | Rosuvastatin (0.5 μM) | 134 | 30.1 |
| OATP1B3 | Rosuvastatin (0.5 μM) | 202 | 51.6 |
| OCT1 | [<sup>14</sup>C]Metformin (10 μM) | >300 | 36.3 |
| OAT1 | [<sup>3</sup>H]PAH (0.5 μM) | >300 | >300 |
| OAT3 | [<sup>3</sup>H]ES (0.1 μM) | >300 | >300 |
| OCT2 | [<sup>14</sup>C]Metformin (20 μM) | >300 | >300 |
| MATE1 | [<sup>14</sup>C]Metformin (20 μM) | >300 | 175.1 |
| MATE2K | [<sup>14</sup>C]Metformin (20 μM) | >300 | 179.8 |

BCRP = Breast cancer resistance protein; BSEP = Bile salt export pump; OAT = Organic anion transporter; OATP = Organic anion-transporting polypeptide; OCT = Organic cation transporter; IC<sub>50</sub> = 50% inhibitory concentration; MATE = Multidrug and toxin extrusion protein; MDR = Multidrug resistance protein; PAH = P-aminohippuric acid; P-gp = P-glycoprotein.

**Table S7.** Plasma protein binding of PF-07304814 or PF-00835231 in plasma, liver microsomes and S9.

| Species | Matrix | Unbound fraction fu (%CV) |  |
| --- | --- | --- | --- |
|  |  | PF-07304814<sup>d</sup> | PF-00835231 |
| Human | Liver microsomes<sup>a,c</sup> | ND | 0.75 (8.7) |
| Human | Liver S9<sup>b,c</sup> | 0.890 (1.9) | ND |
| Human | Plasma | 0.184 (5.9) | 0.449 (7.5) |
| Monkey | Plasma | 0.361 (10.2) | 0.441 (7.7) |
| Dog | Plasma | 0.312 (10.9) | 0.416 (9.5) |
| Rat | Plasma | 0.379<sup>f</sup> (12.7) | 0.327<sup>g</sup> (12.1) |

<sup>a</sup>measured at 0.8mg/mL protein, <sup>b</sup>measured at 0.03mg/mL protein, <sup>c</sup>2 μM substrate concentration, <sup>d</sup>3.4 μM substrate concentration, <sup>e</sup>5 μM substrate concentration, <sup>f</sup>Sprague-Dawley rat plasma, <sup>g</sup>Wistar Hannover rat plasma

**Table S8.**
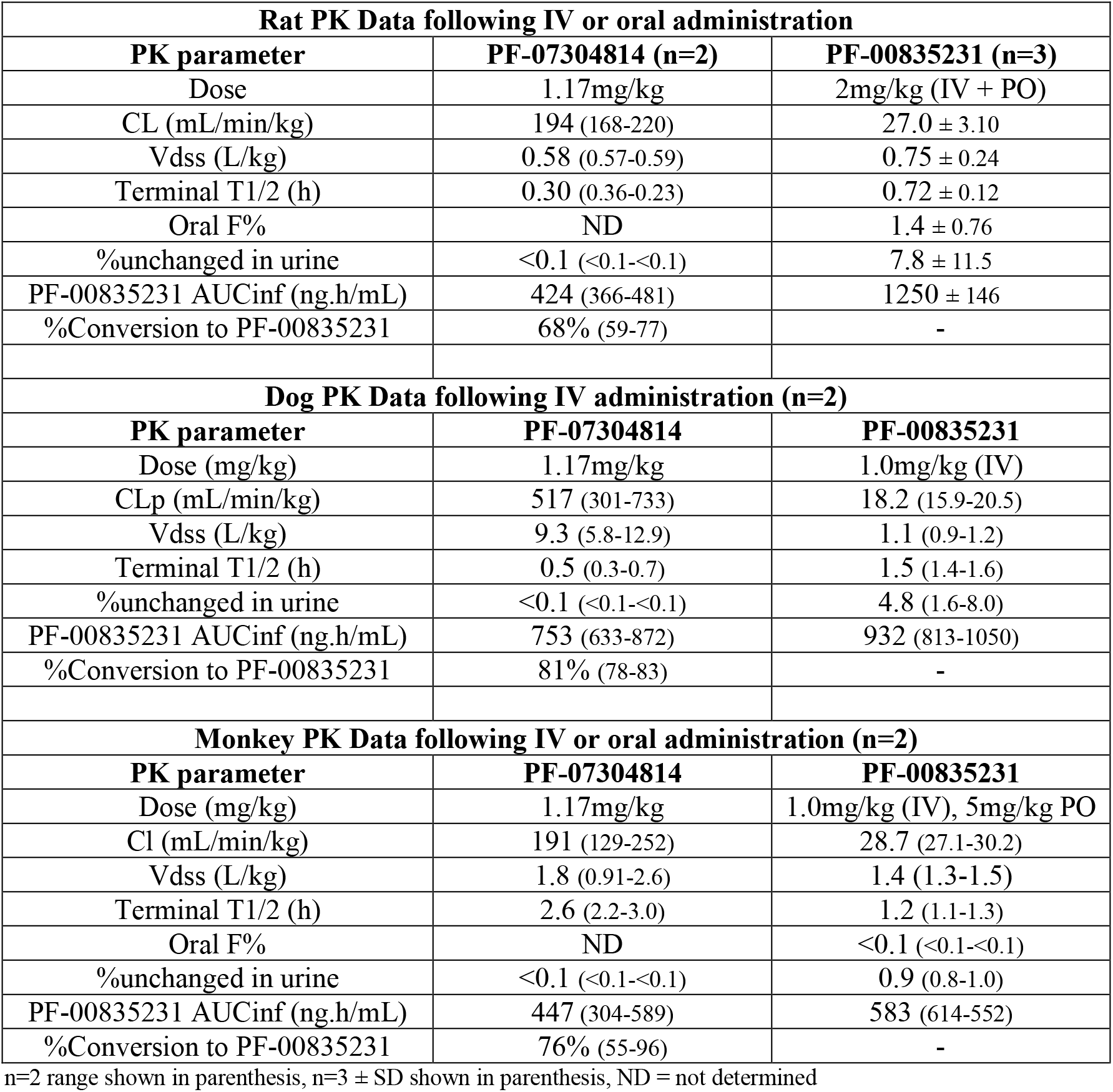
Preclinical plasma PK summary of PF-07304814 or PF-00835231 following oral and/or IV administration to rats, dogs and monkeys.

| **Rat PK Data following IV or oral administration** |  |  |
| --- | --- | --- |
| **PK parameter** | **PF-07304814 (n=2)** | **PF-00835231 (n=3)** |
| Dose | 1.17mg/kg | 2mg/kg (IV + PO) |
| CL (mL/min/kg) | 194 (168-220) | 27.0 ± 3.10 |
| Vdss (L/kg) | 0.58 (0.57-0.59) | 0.75 ± 0.24 |
| Terminal T1/2 (h) | 0.30 (0.36-0.23) | 0.72 ± 0.12 |
| Oral F% | ND | 1.4 ± 0.76 |
| %unchanged in urine | <0.1 (<0.1-<0.1) | 7.8 ± 11.5 |
| PF-00835231 AUCinf (ng.h/mL) | 424 (366-481) | 1250 ± 146 |
| %Conversion to PF-00835231 | 68% (59-77) | - |
| **Dog PK Data following IV administration (n=2)** |  |  |
| **PK parameter** | **PF-07304814** | **PF-00835231** |
| Dose (mg/kg) | 1.17mg/kg | 1.0mg/kg (IV) |
| CLp (mL/min/kg) | 517 (301-733) | 18.2 (15.9-20.5) |
| Vdss (L/kg) | 9.3 (5.8-12.9) | 1.1 (0.9-1.2) |
| Terminal T1/2 (h) | 0.5 (0.3-0.7) | 1.5 (1.4-1.6) |
| %unchanged in urine | <0.1 (<0.1-<0.1) | 4.8 (1.6-8.0) |
| PF-00835231 AUCinf (ng.h/mL) | 753 (633-872) | 932 (813-1050) |
| %Conversion to PF-00835231 | 81% (78-83) | - |
| **Monkey PK Data following IV or oral administration (n=2)** |  |  |
| **PK parameter** | **PF-07304814** | **PF-00835231** |
| Dose (mg/kg) | 1.17mg/kg | 1.0mg/kg (IV), 5mg/kg PO |
| Cl (mL/min/kg) | 191 (129-252) | 28.7 (27.1-30.2) |
| Vdss (L/kg) | 1.8 (0.91-2.6) | 1.4 (1.3-1.5) |
| Terminal T1/2 (h) | 2.6 (2.2-3.0) | 1.2 (1.1-1.3) |
| Oral F% | ND | <0.1 (<0.1-<0.1) |
| %unchanged in urine | <0.1 (<0.1-<0.1) | 0.9 (0.8-1.0) |
| PF-00835231 AUCinf (ng.h/mL) | 447 (304-589) | 583 (614-552) |
| %Conversion to PF-00835231 | 76% (55-96) | - |

n=2 range shown in parenthesis, n=3 ± SD shown in parenthesis, ND = not determined

**Table S9.**
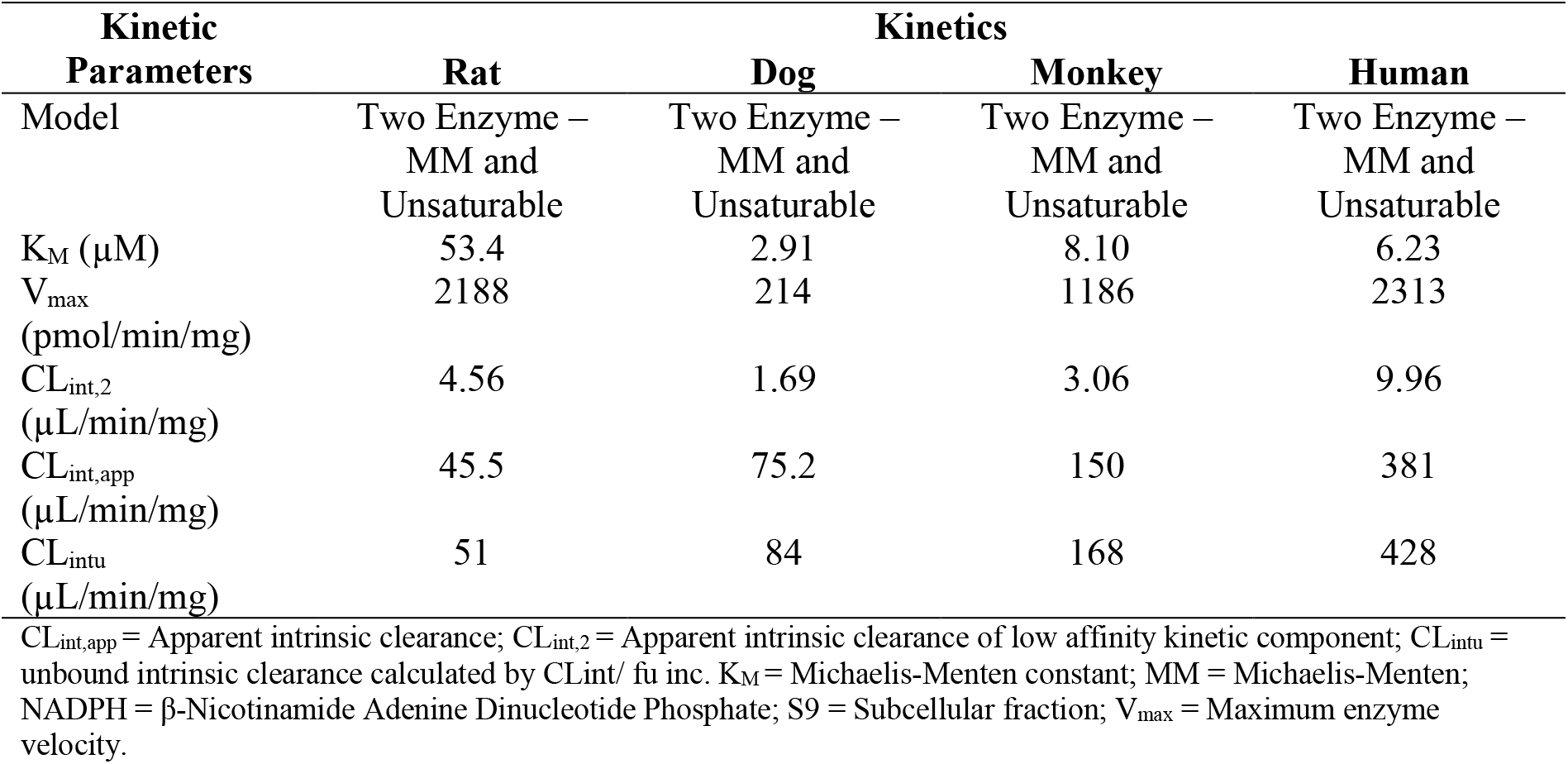
Metabolic stability and enzyme kinetics of PF-07304814 in liver S9.

**Table S10.**
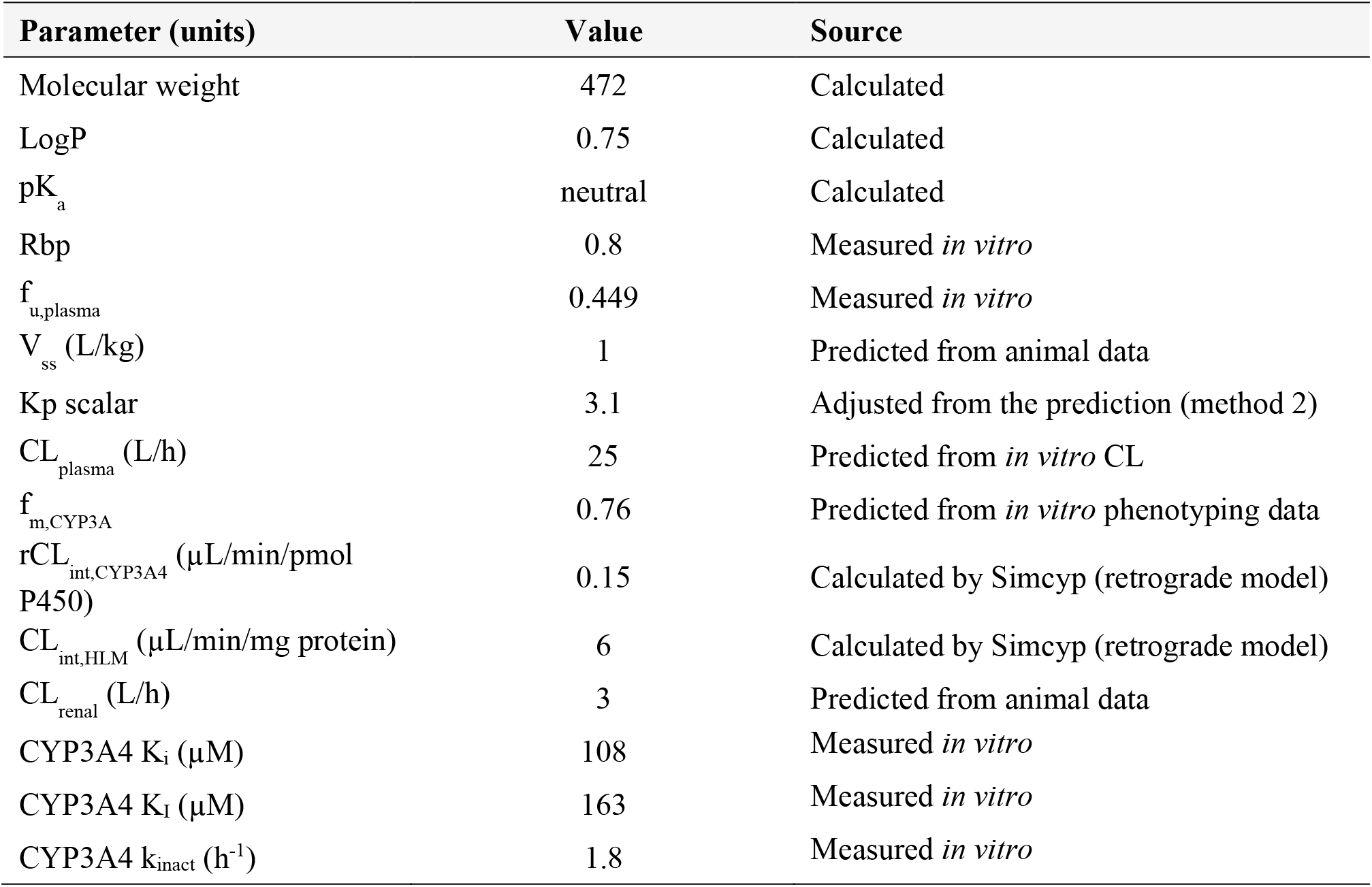
Physicochemical and pharmacokinetic parameters of PF00835231 for PBPK modeling.

**Table S11.** Predicted pharmacokinetic parameters of PF00835231 with and without coadministration of itraconazole.

| PF-07304814<sup>a</sup><br>mg/day | Itraconazole<br>mg/day | C<sub>max</sub> |  | AUC |  | C<sub>max</sub>R | AUCR |
| --- | --- | --- | --- | --- | --- | --- | --- |
|  |  | μM total | μM unbound | μM·h total | μM·h unbound | ratio | ratio |
| 500 | 0 | 1.1 | 0.50 | 27 | 12 | 2.1 | 2.1 |
|  | 200 | 2.4 | 1.1 | 58 | 26 | (2.1-2.3) | (2.0-2.2) |

Data are expressed as geometric mean for C<sub>max</sub> and AUC and geometric mean with 90% confidence intervals in parentheses for C<sub>max</sub>R and AUCR.
<sup>a</sup> Daily dose of PF-07304814 provides formation of PF00835231 equivalent to 320mg/day used in the simulation.

**Table S12.** Human and HIV Protease and Substrate concentrations.

| **Protease/class** | **Enzyme (nM)** | **Substrate** | **Substrate (μM)** |
| --- | --- | --- | --- |
| Caspase 2/ cysteine | 10 | Ac-LEHD-AMC | 5 |
| Cathepsin B/ aspartyl | 1.2 | CBZ-Arg-Arg-AMC | 15 |
| Cathepsin D/ aspartyl | 1.0 | MCA-PLGL-Dap(Dnp)-AR-NH<sub>2</sub> | 2 |
| Chymotrypsin/ serine | 0.5 | Suc-AAPF-AMC | 10 |
| Elastase/ serine | 0.6 | MeOSuc-AAPV-AMC | 10 |
| HIV-1/ aspartyl | 20 | Anaspec SensoLyte | 10 |
| Thrombin a/ serine | 0.01 | H-D-CHA-Ala-Arg-AMC.2AcOH | 10 |

